# The formation of microbial exoskeletons is driven by a controlled calcium-concentrating subcellular niche

**DOI:** 10.1101/2020.01.08.898569

**Authors:** Alona Keren-Paz, Malena Cohen-Cymberknoh, Dror Kolodkin-Gal, Shani Peretz, Iris Karunker, Sharon G. Wolf, Tsviya Olender, Sergey Kapishnikov, Vlad Brumfield, Simon Dersch, Elena Kartvelishvily, Peninnah Green-Zelinger, Damilola Isola-Adeyanju, Ronit Suissa, Michal Shteinberg, Daniel McLeod, Marianna Patrauchan, Gideon Zamir, Assaf Gal, Peter L. Graumann, Eitan Kerem, Ilana Kolodkin-Gal

## Abstract

In nature, bacteria reside in biofilms - multicellular differentiated communities held together by extracellular matrix. In this work, we identified a novel subpopulation essential for biofilm formation – mineral-forming cells. This subpopulation contains an intracellular calcium-accumulating niche, in which the formation of a calcium carbonate mineral is initiated. As the biofilm colony develops, this mineral grows in a controlled manner, forming a functional macrostructure that serves the entire community.

The molecular mechanisms promoting calcite scaffold formation were conserved between three distant phyla – the Gram-positive *Bacillus subtilis*, Gram-negative *Pseudomonas aeruginosa* and the actinobacterium *Mycobacterium abscessus*. Biofilm development of all three species was similarly impaired by inhibition of calcium uptake and carbonate accumulation. Moreover, chemical inhibition and mutations targeting mineralization both significantly reduced the attachment of *P. aeruginosa* to the lung, as well as the subsequent damage inflicted by biofilms to lung tissues, and restored their sensitivity to antibiotics.

The evolutionary conserved cellular pathway controlling the fundamental feature of biofilm development uncovered in this work offers novel druggable targets for antibiotics to combat otherwise untreatable biofilm infections.

## Introduction

While bacteria have been historically studied in homogeneous monocultures, in natural ecosystems and in clinical settings they are typically found in interface-associated multicellular communities called biofilms (Kolter and Greenberg, 2006). It was recently indicated that in soils, oceans, the deep subsurface, phyloshpere and animal microbiomes, 40−80% of bacteria reside in biofilms (Flemming and Wuertz, 2019). Bacterial biofilms are differentiated communities, in which the cells are held together by extracellular matrix. The formation of a biofilm is frequently favored over free-living planktonic life-style, as it increases the fitness of member cells promoting their survival, such as better attachment to hosts, division of metabolic labor and protection from the environment.

Microbial biofilms are of extreme clinical importance, as they are associated with many persistent and chronic infections (Costerton et al., 1999). A prominent feature of biofilms is their inherent resistance to the immune system and to antibiotics (Bryers, 2008; Bucher et al., 2019; Hill et al., 2005) – biofilm cells are up to 1,000 times more tolerant than planktonic bacteria (Bryers, 2008), making the eradication of biofilm infections with currently available antibiotics extremely challenging.

For example, the commensal/aquatic bacterium *Pseudomonas aeruginosa* can cause devastating chronic biofilm infections in compromised hosts, such as cystic fibrosis (CF) patients and those with burn wounds or implanted medical devices (Costerton et al., 1999). Respiratory infections with *P. aeruginosa* are a leading cause of morbidity and mortality in patients with CF (Yoon and Hassett, 2004). Once a chronic infection with *P. aeruginosa* is established, it is almost impossible to eradicate (Cohen-Cymberknoh et al., 2016). Similarly, the biofilms of *Mycobacterium abscessus* cause infections of respiratory tract, skin and central nervous system in patients with CF and chronic obstructive pulmonary disease (Caimmi et al., 2018).

A hallmark of biofilms is their ability to form robust and complex 3D structures, and the distinctive spatial organization provides the residing bacteria with several benefits. The biofilm architecture was suggested to relieve metabolic stress. For example, channels formed below the ridges and wrinkles within the colony may facilitate diffusion of fluids, nutrients and oxygen (Bloom-Ackermann et al., 2016; Dietrich et al., 2013; Kolodkin-Gal et al., 2013; Wilking et al., 2013). Furthermore, cells located in different areas of the colony are exposed to different levels of oxygen, nutrients and quorum sensing molecules, which affect the genetic programs they express, promoting differentiation (Asally et al., 2012; Hassanov et al., 2018; Liu et al., 2015; Monds and O’Toole, 2009; Serra and Hengge, 2014; Stewart and Franklin, 2008). Thus, community structure allows the genetically identical biofilm cells to display phenotypic heterogeneity and to endure stressful environments. Due to physical protection offered by the biofilm biomass, differentiation and physiological adaptations, cells residing deep within the biofilm are protected from environmental assaults, such as antibiotics. Biofilm colonies are a compelling model system for biofilm development, and many basic biological processes of high relevance to bacterial pathogenicity were first discovered in this well-controlled and robust system (Colvin et al., 2012; Dietrich et al., 2008; El Mammeri et al., 2019; Hufnagel et al., 2018; Jo et al., 2017; Richards et al., 2019; Serra et al., 2013; Steinberg et al., 2020; Wermser and Lopez, 2018).

Until recently, the ability of biofilm-forming bacteria to generate complex architectures was attributed exclusively to self-produced **organic** extracellular matrix (ECM) (Dragos and Kovacs, 2017; Reichhardt and Parsek, 2019; Steinberg and Kolodkin-Gal, 2015), composed of carbohydrate-rich polymers (i.e., lipids or exopolysaccharides), proteins, and nucleic acids (Branda et al., 2005). However, we and others have recently shown that microbial biofilms contain an organized internal mineral structure, composed of crystalline calcium carbonate (calcite), that also contributes to their 3D morphology (Keren-Paz et al., 2018; Keren-Paz and Kolodkin-Gal, 2020; Li et al., 2015).

Formation of minerals is known to be induced in bacterial biofilms, through passive surface-mediated processes promoted by by-products of bacterial metabolism. Biologically induced minerals associated with bacteria include oxides of Fe, Mn, and other metals; metal sulfates and sulfites; phosphates and carbonates; and Fe and Fe-Al silicates (Frankel and Bazylinski, 2003). Of all examples of biomineralization, microbial induced calcium carbonate precipitation (MICCP) is most frequently associated with microbial communities (Weiner and Dove, 2003), (Douglas and Beveridge, 1998). In geological settings, bacterial metabolic processes, such as urea hydrolysis by urease-positive bacteria, increase local concentration of bicarbonate and elevate pH. When enough environmental calcium is present, those changes promote spontaneous precipitation of calcium carbonate, with bacterial envelopes serving as nucleation sites for the growing mineral, leading to formation of crystalline calcium carbonate (Dhami et al., 2013). Similar processes were also suggested to promote biofilm-associated calcification of *Proteus mirabilis* on catheters (Morris and Stickler, 1998), and during dual-species biofilm formation (Li et al., 2016b) and to contribute to *P. aeruginosa* virulence in an insect host (Lotlikar et al., 2019). In both clinical and environmental scenarios, calcium deposition in biofilms was mostly seen as an uncontrolled and unintentional byproduct of bacterial metabolic activity.

Recently, intracellular amorphous calcium carbonate (ACC) granules were detected in some species of Gram-negative autotropic bacteria (Blondeau et al., 2018). Those intracellular deposits were suggested to contribute to single cell energy metabolism by promoting photosynthesis (Blondeau et al., 2018), or chemolithoautotrophy (Monteil et al., 2020). While the precise molecular mechanisms regulating this process are still unresolved, the presence of a microcompartment dedicated to mineralization raises the possibility that some bacteria might be capable of controlling the formation of biogenic mineral.

Our recent discovery of precisely organized mineral macrostructures within biofilms suggests that biomineralization in biofilms is tightly controlled. These calcite ‘skeletons’ contributed to fitness of biofilm colonies in two unrelated soil bacteria: *Bacillus subtilis* and *Mycobacterium smegmatis* (Keren-Paz et al., 2018; Oppenheimer-Shaanan et al., 2016). For both, the mineral structure acted as a structural scaffold supporting the 3D architecture of the colony, and as diffusion barrier preventing the penetration of solutes into the biofilms. The formation of controlled calcium carbonate macrostructures in phylogenetically distinct heterotrophic biofilms raises the possibility that the function of calcium carbonate is not limited to specific species utilizing it for intracellular chemical energy, but is instead a general phenomenon wide-spread in the bacterial kingdom.

Those reports challenge the current view of biofilm development as a process depending solely on organic ECM production. They raise several fundamental questions regarding the developmental role of mineral skeleton formation within a bacterial community, and the relation between structure and function in microbial biofilms. If indeed mineralization is a controlled process resulting in a structure crucial to biofilm development and function, this is a novel aspect of basic biofilm biology. What is the role of mineral structure in the development of differentiated biofilm community? Are there dedicated ‘osteoblast-like’ cells initiating mineral production? What are the fitness advantages that the mineral exoskeleton confers to the community? Furthermore, organic matrices and their regulation differ between different organisms (Steinberg and Kolodkin-Gal, 2015). However, the formation of calcium carbonate relies on a highly conserved building blocks, calcium and carbonate, generated from carbon dioxide in most heterotopic organisms. Therefore, the generation of mineral scaffolds could be common and conserved across the bacterial domain, contributing to the phenotypical resistance of diverse microbial biofilms. If so, understanding the underlying molecular mechanisms of biofilm mineralization is a crucial step towards our ability to successfully combat biofilm infections.

In this work, we discovered that the calcium-dependent 3D organization of the biofilm colony leads to transcriptional reprogramming of the bacterial community, and was essential for biofilm development. Calcite formation and the assembly of a functional mineralized macro-skeleton supporting the 3D morphology of a biofilm colony was associated with defined intracellular calcium-rich deposits present in a distinct subpopulation of biofilm cells, differing from flagellin and organic ECM producers. Calcite formation turns out to be an extremely conserved process, occurring not only in the beneficial model organism *B. subtilis*, but also in two unrelated lung pathogens – *P. aeruginosa* and *M. abscessus*. Inhibition of key biomineralization enzymes or calcium uptake prevented biofilm formation by both pathogens. We were able to identify calcite in sputum samples taken from CF patients, suggesting that this conserved process is of clinical importance. Finally, in an *ex vivo* lung model, chemical and genetic inhibition of calcium uptake and of carbonate accumulation blocked biofilm formation and lung colonization, preventing damage inflicted by *P. aeruginosa* to lung tissues, and sensitized *P. aeruginosa* to antibiotic treatment.

Taken together, our results identify a previously overlooked process essential for bacterial biofilm development – **the tightly regulated formation of mineral skeletons by dedicated cells**. Its conservation across the bacterial kingdom highlights the fundamental role it plays in biofilm biology, and could lead to novel therapeutic approaches for combating a broad range of persistent biofilm infections.

## Results

In a previous work, we demonstrated that *B. subtilis* biofilm morphology is calcium-dependent (Oppenheimer-Shaanan et al., 2016). Removing calcium acetate from the undefined biofilm-promoting B4 medium prevented the development of the complex 3D architecture characteristic of a biofilm colony, without affecting bacterial growth. This 3D structure required calcium (as no equivalent cation could replace it and as it was induced by various calcium salts) and metabolically produced carbonate (as it was eliminated under anaerobic conditions and increased in CO_2_-enriched environment). The mineral component was identified as calcite by FTIR and XRD (Supplementary Fig. 1, (Oppenheimer-Shaanan et al., 2016). It spanned the entire colony and was spatially co-localized with colony wrinkles, suggesting it acted as a “scaffold” supporting them (Oppenheimer-Shaanan et al., 2016). The mineral structure developed and increased over time as the colony grew (Keren-Paz et al., 2018). Finally, as the biofilm colonies aged and their 3D spatial structure diminished, the internal and organized mineral deteriorated (potentially due to fermentation and acidification of the medium by oxygen-depleted cells within the inner mass of the aging biofilm), and only large non-structural crystals were left in the colony periphery due to a passive growth of calcite crystals (Oppenheimer-Shaanan et al., 2016). During biomineralization, the formation of exopolysaccharides and the amyloid protein TasA was also essential to form complex morphology, and interactions between the organic and inorganic matrix were evident (Oppenheimer-Shaanan et al., 2016).

To better understand the molecular mechanisms underlying the formation of functional mineral macrostructures, we examined the effect of calcium on gene expression in a *B. subtilis* biofilm colony by sequencing the transcriptome of colonies grown either with or without excess calcium (Fig. 1A). The addition of calcium resulted in a dramatic reprogramming of the transcriptional profile of the colony, with 20% of the genome (n=875) significantly changing between the two conditions at all time-points tested (Supplementary File 2). Only in the presence of added calcium, specific developmental processes defining the biofilm state, such as induction of ECM production and sporulation, and the repression of motility, were observed (Fig. 1B). These fundamental changes were mostly sustained over time (Fig. 1C). We next compared the two transcriptional signatures to a collection of published data, containing 269 mRNA profiles of *B. subtilis* grown under over 100 different conditions (Nicolas et al., 2012) (Fig. 1D). The transcriptome of cells grown with excess calcium was similar to that of previously analyzed biofilm cells, regardless of the biofilm promoting medium used. On the other hand, the transcription profile of colonies grown without added calcium was similar to that of planktonic cells experiencing nutrient starvation, but was not identified as biofilm in this unbiased analysis.

**Figure 1:**
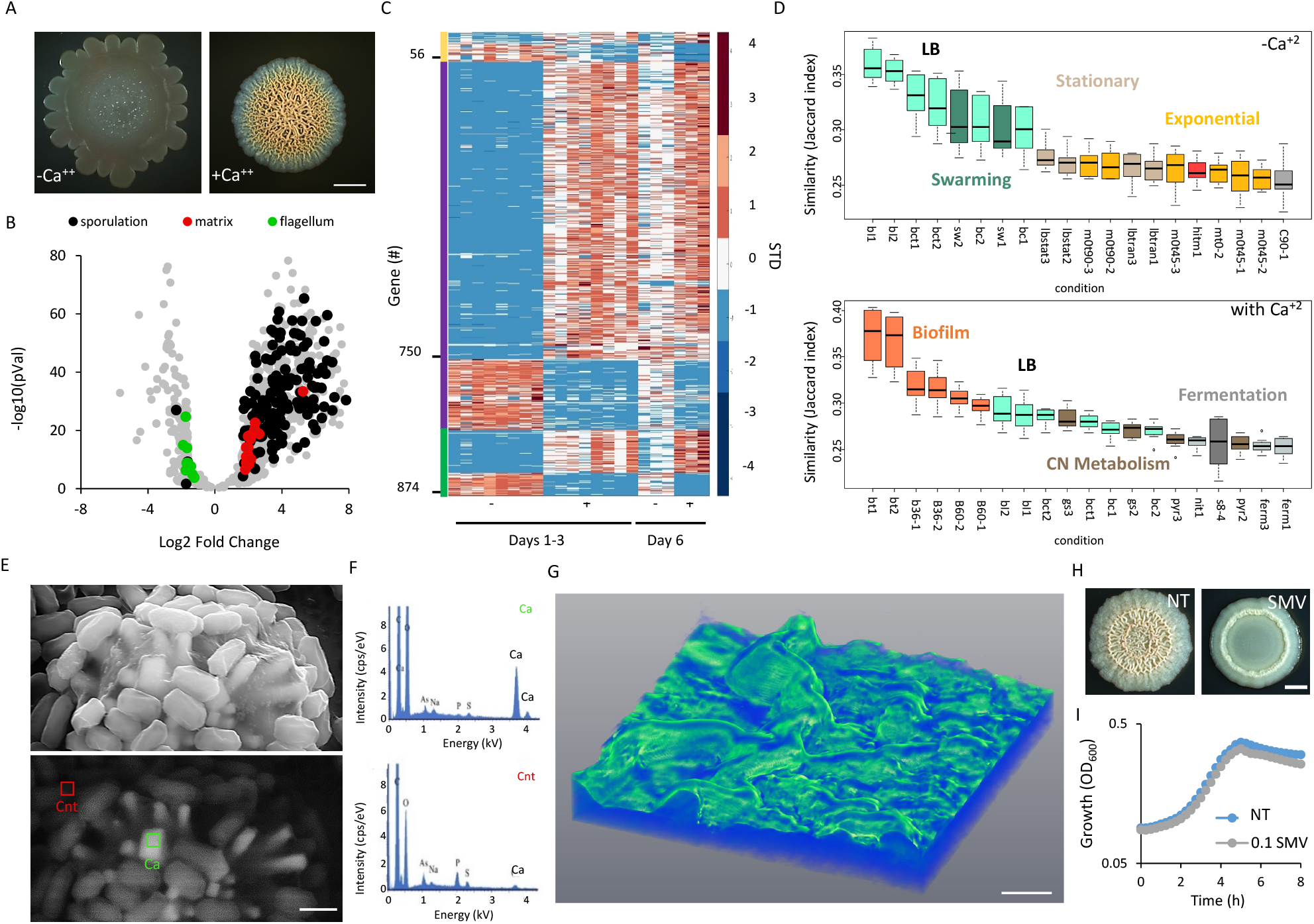
**Calcium is necessary for normal biofilm development and structure** A. Light microscopy images of 6-day-old *B. subtilis* NCIB 3610 biofilm colonies grown on B4 agar, showing the effect of addition of calcium (0.25% v/v calcium acetate) on colony architecture. Scale bar – 2 mm. A representative image (out of n = 3 experiments) is shown. B. Volcano-plot depicting calcium-dependent changes in the transcriptome in *B. subtilis* biofilm colonies. Significance (P values) vs difference (fold changes) are plotted. Functional categories were determined by DAVID analysis. C. A heatmap with all differentially expressed genes (n = 876), scaled to the mean expression level of each gene. The map is ordered by age of biofilm colonies, as shown by the left colored bar, where green are genes that are differentially expressed in days 1-3, and day 6; purple genes that are differentially expressed in days 1-3 and yellow genes that are differentially expressed in day 6. D. The top 20 conditions (Nicolas et al., 2012), showing the highest similarity (expressed as Jaccard index) to the transcriptome of *B. subtilis* 6-day-old biofilm colonies grown with or without calcium. E. Biofilm cells closely associated with prismatic mineral structures, as visualized by Scanning Electron Microscopy (SEM) image of 10-day-old *B. subtilis* biofilm colony grown with calcium. Upper panel - secondary mode; lower panel - backscattering mode. Magnification – X25000, scale bar – 1 µm. A representative field (out of n=5 fields, from 6 experiments) is shown. F. EDX analysis of areas indicated in (E), indicating that mineral structures are rich in calcium. Ca – calcium-rich, Cnt – control. G. 3D re-construction of microCT X-Ray analysis revealing 3D mineral distribution in a 5-day old *B. subtilis* colony grown with calcium. Color indicates intensity, with green representing the densest mineral. Scale bar – 0.2 mm. A representative image (out of n = 3 experiments) is shown. H. Light microscopy images of 5-day-old *B. subtilis* biofilm colonies grown with calcium, and supplemented with calcium uptake inhibitor (0.1 mg/ml sodium metavanadate (SMV)), as indicated. NT indicated untreated culture. Scale bar – 2 mm. A representative image (out of n = 3 experiments) is shown. I. The effect of inhibiting calcium uptake on planktonic growth. *B. subtilis* was grown shaking in liquid LB medium with calcium, supplemented with 0.1 mg/ml SMV, as indicated. Results are averages of nine wells, bars represent standard deviations. A representative experiment (out of n=3 experiments) is shown.

These findings highlight the central role of calcium in biofilm development, and suggest an intimate connection between biofilm structure and function - raising the possibility that microbial cells are actively regulating calcium carbonate biomineralization. Indeed, when a biofilm colony was visualized by scanning electron microscopy (SEM), we observed a highly mineralized subpopulation of cells, tightly associated with mineral crystals (Fig. 1E, Supplementary Fig. 2). Consistently with our previous characterization of bleached biogenic calcite, the unbleached crystals had rough faces, along with a smooth and flat crystal face and displayed elongated prismatic morphology instead of the rhombohedral morphology of calcite grown in pure solution (Oppenheimer-Shaanan et al., 2016).

The crystals were consistent with biogenic calcite generated by *B. subtilis (Oppenheimer-Shaanan et al., 2016)*, as judged by shape; backscatter mode and energy dispersive X-ray spectroscopy (EDX) (Fig. 1F, Supplementary Fig. 2). In the absence of added calcium, mineral crystals were rarely detected (Supplementary Fig. 3). Mineral producers were not associated with organic ECM, did not carry flagella, and did not sporulate (Figures 1E, Supporting Figures 2 and 3), and therefore morphologically differed from previous subpopulations observed in *B. subtilis* biofilms (Vlamakis et al., 2008). To create the 3D reconstruction, a whole, unfixed bacterial colony was transferred to a plastic slide for a microCT scan, and rotated between the X-ray source and the detector positioned at optimal distances for a voxel size of 0.87 µm. 2D projections were taken at different angles until a full rotation (360^°^) was completed. The full set of images was then used to reconstruct the whole volume of the sample by back projection algorithm and thus a high-resolution 3D image was generated (see Supplementary Movie 1). This microCT X-ray scan confirmed that calcite is organized in a macro-structure, generating a non-uniform continuous layer throughout the wrinkles (Fig. 1G).

**Figure 2:**
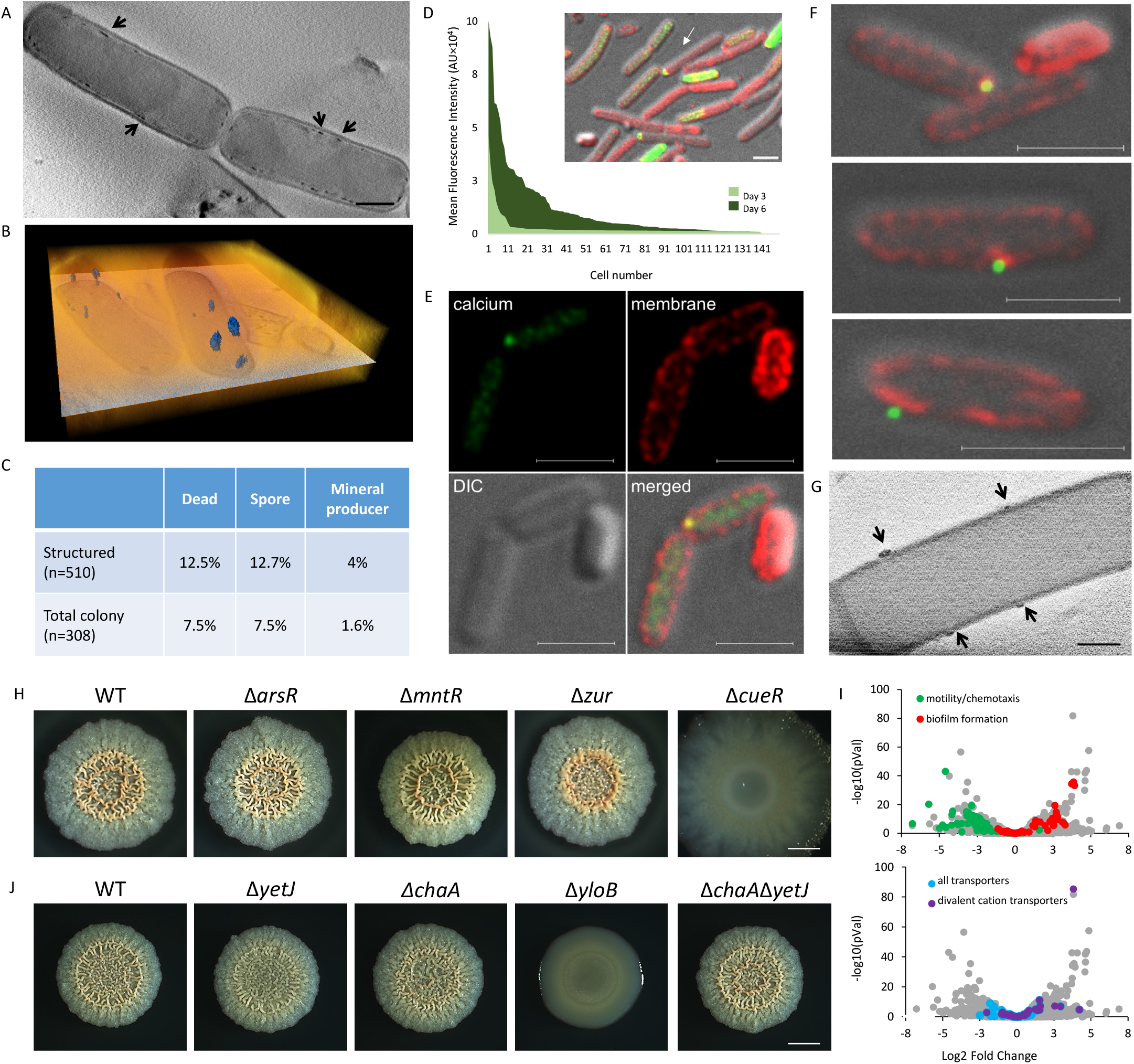
**Mineralization is initiated intracellularly** A. A 30-nm thick virtual slice through a CSTET 3D reconstruction of *B. subtilis* cells showing intracellular calcium-rich deposits (arrows). Scale bar – 400 nm. A representative field (out of n=20 fields, from 3 experiments) is shown. B. Volume rendering (orange) and a single orthoslice (greyscale) through the center of the volume, from a 3D CSTET reconstruction. The calcium-rich deposits are artificially colored (blue). C. Quantification of different subpopulations within a 6-day old *B. subtilis* biofilm colony, showing enrichment of cells with mineral foci in the structured areas (wrinkles) of the colony. D. Intracellular calcium levels of bacterial cells (n=150) stained with calcein-AM, a calcium-specific fluorescent dye. Cells were isolated from *B. subtilis* biofilm colonies (day 3 and 6) grown with calcium. Inset – a representative image of stained cells isolated from a 6 day-old biofilm. Green – calcein-AM, red – NileRed membrane stain, gray – DIC. Scale bar – 2 µm. E. and F. G-STED images *B. subtilis* cells isolated from 6 day-old biofilm colonies grown with calcium. Green – calcein-AM, red – NileRed membrane stain, gray – DIC. Scale bar – 2 µm. G. 30-nm thick virtual slice through a CSTET 3D reconstruction. Arrows indicate calcium-rich deposits associated with a membrane from the outside of the cell. Scale bar – 400 nm. H. Light microscopy images of 3-day-old biofilm colonies of wild-type *B. subtilis* and its mutant derivatives (see text for details) grown with calcium. Scale bar – 2 mm. A representative experiment (out of n=3 experiments) is shown. I. Volcano-plot depicting CueR-dependent changes in expression of genes in 3-day old *B. subtilis* biofilm colony. Significance (P values) vs difference (fold changes) are plotted. Functional categories were determined by DAVID analysis. J. Light microscopy images of 2-day-old biofilm colonies of wild-type *B. subtilis* and its mutant derivatives (see text for details) grown with calcium. Scale bar – 2 mm. A representative experiment (out of n=3 experiments) is shown.

**Figure 3:**
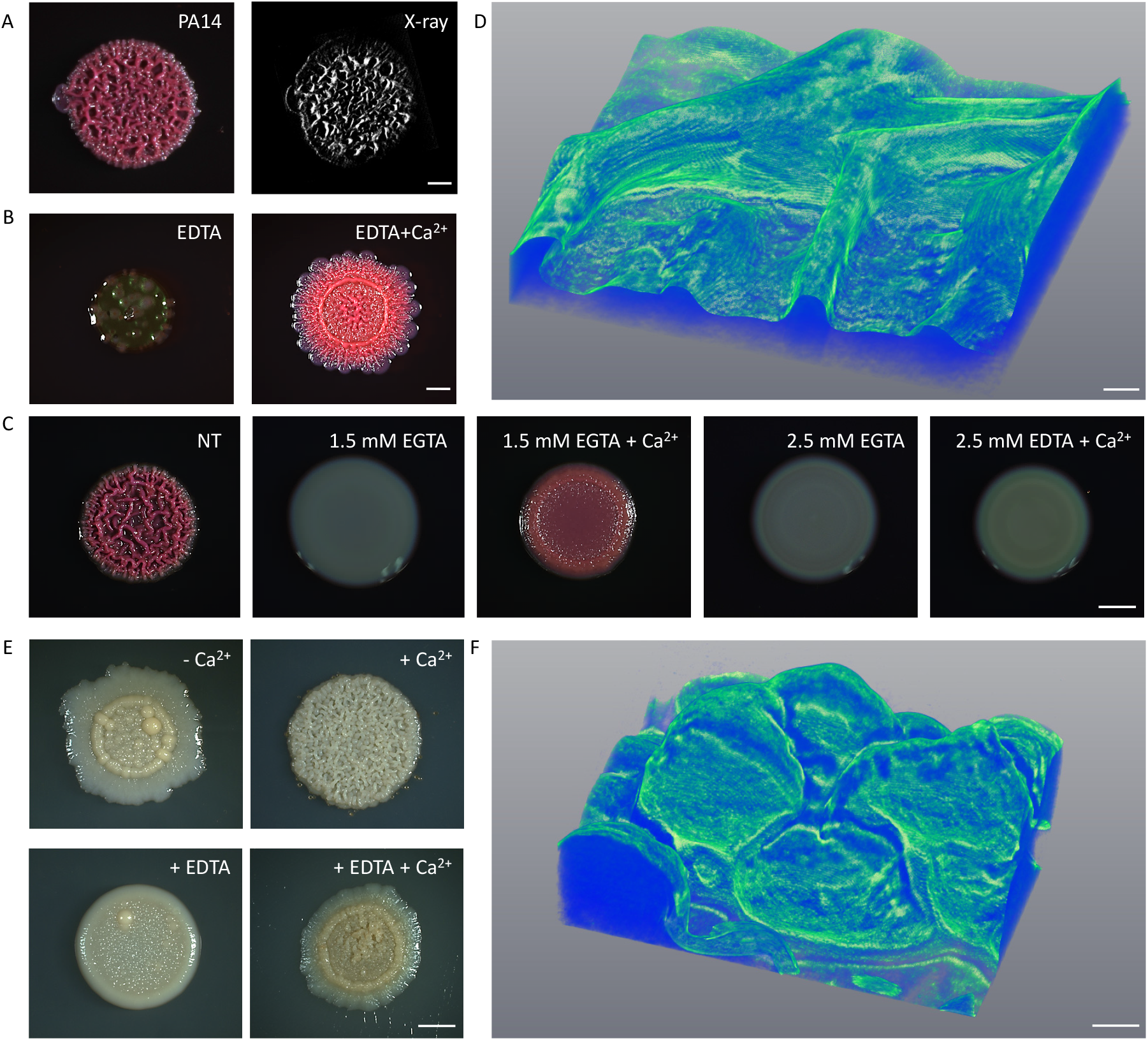
**The role of calcium in the biofilm structure is conserved across bacterial species** A. Light microscopy (left) and microCT-X-ray (right) images of 3-day-old *P. aeruginosa* PA14 biofilm colonies grown on TB agar. In the X-Ray images, the bright contrast appears white, and indicates the location of dense mineral, while the organic matter appears dark. Scale bar – 1 mm. A representative image (out of n = 3 experiments) is shown. B. Light microscopy images of 3-day-old *P. aeruginosa* PA14 of biofilm colonies grown on TB agar supplemented with 1 mg/ml EDTA (chelating agent), either with or without the addition of 700 mM CaCl_2_. Scale bar – 1 mm. A representative experiment (out of n=3 experiments) is shown. C. Light microscopy images of 3-day-old *P. aeruginosa* PA14 biofilm colonies, grown on TB agar either untreated (NT), or supplemented with 700 mM CaCl_2_ and EGTA (calcium chelator), as indicated. Scale bar – 5 mm. A representative image (out of n = 3 experiments) is shown. D. 3D re-construction of microCT X-Ray analysis revealing mineral distribution in a 5-day old *P. aeruginosa* PA14 colony. Color indicates intensity, with green representing the densest mineral. Scale bar – 0.2 mm. A representative image (out of n = 3 experiments) is shown. E. Light microscopy images of 5 day-old *M. abscessus* biofilm colonies, grown on B4 agar, supplemented with either calcium acetate (0.25% v/v), or 1.5 mM EDTA, as indicated. Scale bar – 5 mm. A representative image (out of n = 3 experiments) is shown. F. 3D re-construction of microCT X-Ray analysis revealing mineral distribution in a 5-day old *M. abscessus* colony grown on B4 agar with calcium. Color indicates intensity, with green representing the densest mineral. Scale bar – 0.1 mm. A representative image (out of n = 3 experiments) is shown.

One possible mechanism linking calcium mineralization and colony structure is that, in the presence of calcium, some kind of specialized cells located in specific regions of the colony, would serve as nucleation sites and promote localized mineral formation. The observed induction of sporulation regulon transcription by calcium (Fig. 1B) raised the possibility that spores could serve this function, possibly due to unique membrane properties of the spore. However, a mutant strain lacking sporulation sigma factor SigF (de Hoon et al., 2010), which does not form spores and is blocked early during sporulation, was able to form highly structured colonies, in a calcium-dependent manner (Supplementary Fig. 4), suggesting that biomineralization promotes biofilm development and sporulation, and not *vice versa*.

The tight association of calcium carbonate mineral with the bacterial cells led us to examine the role of intracellular calcium. Disruption of intracellular calcium homeostasis by a chemical inhibitor of P-type ATPases sodium metavanadate (SMV) (Clausen et al., 2016; Guragain et al., 2013; Kuhlbrandt, 2004), prevented the formation of a complex 3D structure even in the presence of excess calcium (Fig. 1H). This morphological defect was not due to inhibition of planktonic growth, which was only mildly affected by SMV (Fig. 1I). This need for controlled intracellular calcium levels raised the possibility that carbonate mineralization starts intracellularly. Therefore, we next examined single cells from biofilms grown with excess calcium.

3D reconstructions of cryo-fixed bacteria with scanning transmission electron tomography (CSTET) (Wolf et al., 2014) (Fig. 2A-B) revealed calcium dense foci inside the cells, associated with the membrane. CryoSTEM imaging and cryo-EDX measurements verified excess calcium in these deposits (Supplementary Fig. 5). The appearance of these deposits was calcium dependent, as we could not observe them in cells grown without added calcium (Supplementary Fig. 6). While the mineral producing sub-population was small, it was enriched in the structured zones of the colony (Fig. 2C). Next, we examined the distribution of cellular calcium levels within the biofilm community. We stained live biofilm cells with calcein-AM – a dye that can be used for visualization of intracellular calcium in live cells (Hale et al., 2000). This lipophilic stain can freely enter the cells, but only binds calcium and becomes fluorescent after the acetoxymethyl (AM) groups are removed by cellular esterases. Gated-STED fluorescent microscopy revealed heterogeneity in cellular calcium levels within the biofilm colony, with most cells containing low levels of calcium, while a smaller fraction contained very high levels (Fig. 2D). The number of cells showing a strong calcein signal increased during biofilm development, as the calcified structures matured. While the cytoplasm of most cells was stained evenly, a small fraction of cells (2.6% out of 300 cells examined) displayed concrete, membrane-localized calcium deposits (Fig. 2E), comparable in their frequency, distribution and localization with calcium deposits identified by CSTET (Fig. 2A-C). Consistent with our previous observation, sporulation does not seem to be involved in cellular calcium deposition – as Δ*sigF* cells contained calcium-rich foci similar to those of wild type (Supplementary Fig. 7). Furthermore, we could detect such deposits associated with the cell envelope on the cytoplasm side, embedded within the membrane, and closely associated with the cell from the outside – suggesting different stages of export (Fig. 2F). Similar calcium deposits were also observed by CSTET in close association with the outer membrane of bacterial cells (Fig. 2G). Taken together, those finding support the existence of a calcium-concentrating compartment in a subpopulation of heterotropic biofilm cells, similar to that recently discovered in autotrophic bacteria (Blondeau et al., 2018), and that the mechanism for its formation involves an active and regulated calcium uptake, as well as the export of the stored calcium (Fig.2F).

In *B. subtilis*, the transcriptional regulation of calcium uptake is poorly characterized. We therefore examined the potential involvement of several general transcriptional regulators of cation uptake. Non-ferric metal homeostasis in bacteria is regulated by the MntR, MerR and ArsR familes of transcriptional regulators (Moore and Helmann, 2005). Strains lacking *arsR*, *mntR* and *zur* developed robust and structured biofilm colonies. In contrast, *cueR* mutant had a severely altered biofilm phenotype and formed completely flat colonies (Fig. 2H). CSTET analysis was suggestive of diminished calcium-concentrating population (a maximum of 1.3% compared with 4% of the wild type, n=149). CueR is a poorly characterized transcriptional regulator belonging to the metal-responsive MerR family (Moore and Helmann, 2005). CueR binds the promoter of copper transporting P-type ATPase CopA, but a deletion of *copA* had no effect of biofilm morphology (Supplementary Fig. 8). The effect on biofilm morphology was also not due to reduction in growth of *cueR* mutant (Supplementary Fig. 9). Those observations raise the possibility that CueR regulates additional, yet undiscovered, P-type ATPase, involved in calcium uptake. To characterize the regulon of CueR, we repeated the transcriptome analysis of the wild-type strain and of *cueR* mutant, both grown in the presence of calcium. When compared to the wild type colony, the expression of matrix genes was upregulated and motility was inhibited in the *cueR* mutant, suggesting a general dis-regulation of biofilm transcriptional profile (Fig. 2I). Biofilm formation is a tightly regulated developmental process, and both inhibition and overexpression of key components is known to cause severe defects in biofilm development (for example, both deletion and over-activation of the biofilm regulator Spo0A prevent normal biofilm development (Veening et al., 2006). High levels of ECM gene expression further indicates that the featureless morphology of the *cueR* mutant is not a result of the lack of organic ECM components. The subsequent analysis of the expression of divalent cation transporters reveled that their expression is significantly altered in *cueR* mutant, while other transporters are less affected (Fig. 2I). Among those, we identified three calcium transporters with transcriptional levels significantly changed in the mutant: *yloB* (ATP-driven Ca^2+^ pump) (Gupta et al., 2017) *, chaA* (H^+^/Ca^2+^ exchanger) (Fujisawa et al., 2009) and *yetJ* (pH-dependent calcium leak channel) (Chang et al., 2014) (Supplementary File 2). Deletion of *yloB* (but not *chaA*, *yetJ*, or both) was sufficient to prevent 3D colony morphology formation, indicating the importance of calcium uptake and homeostasis (Fig. 2J and Supplementary Fig. 10).

Taken together, these observations suggest that the first critical step in the formation of calcite mineral is initiated in saturated mineral precursor microenvironments within the cell. After the initial nucleation, the precursor seems to be exported out of the bacterial cell without hampering the membrane integrity of its producer, and to further grow on organic ECM templates in the highly complex biofilm microenvironment, until an organized functional mineral macro-structure is formed.

The organic ECM components and the molecular mechanism governing transition from planktonic life style to the biofilm state vary immensely between phylogenetically distant bacteria, making a development of broad range drugs specifically targeting biofilms challenging. In this sense, biomineralization is an extremely appealing target, as it relies on simple and conserved mechanisms allowing accumulation of calcium and carbonate. We have recently reported that, like *B. subtilis*, the biofilm colonies of soil actinobacterium *M. smegmatis* contain functional mineral macro-structures (Oppenheimer-Shaanan et al., 2016). One additional study provided evidence for some spatial organization of calcium carbonate mineral in submerged biofilms formed by the Gram-negative pathogen *P. aeruginosa* (Li et al., 2016a). This led us to explore whether we can detect regulated formation of functional mineral structures in pathogenic biofilm colonies.

When grown on biofilm-inducing agar, *P. aeruginosa* PA14 forms complex colonies with a highly characteristic morphology, which were shown to be dependent on polysaccharides encoded by the *pel* operon (Colvin et al., 2011)). However, like in *B. subtilis,* these complex 3D structures were co-localized with spatially organized mineral, as determined by microCT (Fig. 3A), were abolished by the cation chelator EDTA and EGTA, and could be restored by addition of excess calcium (Fig. 3B, C). 3D reconstruction confirmed that the densest mineral areas formed non-continuous layer correlated with colony wrinkles (Fig. 3D), further suggesting that the 3D structure of *P. aeruginosa* biofilms likely relies on the formation of calcite mineral scaffolds. The formation of elaborate calcium-dependent 3D morphology was conserved in an additional strain of *P. aeruginosa* PA01 (Supplementary Fig. 11). The mineral and exopolysaccharides were both essential for structure formation, as a mutant for both *pel* and *psl* operons was flat and featureless, similarly to biofilms grown in the presence of a calcium chelator (Supplementary Fig. 12).

To indicate whether the formation of mineral scaffolds is a conserved process of significant clinical potential, we tested whether it occurs in another, phylogenetically distant, drug-resistant pathogen. We therefore examined the biofilms of *M. abscessus,* which are held together by extracellular matrix consisting of mycolic acids (Halloum et al., 2016), which bear little resemblance to EPS components of *B. subtilis* and *P. aeruginosa*. Nevertheless, this bacterium also only formed colonies of complex morphology in the presence of calcium (Fig. 3E). Fourier-transform infrared spectroscopy (FTIR) detected calcium carbonate mineral in the colony (Supplementary Fig. 13), and X-Ray analysis revealed that the densest mineral areas were co-localized with colony wrinkles (Fig. 3F), just like in *B. subtilis* and *P. aeruginosa* colonies. In addition to the organized macro-scale structures present throughout the colony, over time large crystals also formed beneath the colony and in its periphery, reminiscent of crystals passively precipitated in the vicinity of *B. subtilis* colonies at later stages. All the bacteria we tested formed mineral macrostructures in a slightly basic-neutral macro pH (Supplementary Fig. 14). Under such conditions, passive mineral precipitation is unlikely, supporting a potential biological regulation of mineralization.

**Figure 4:**
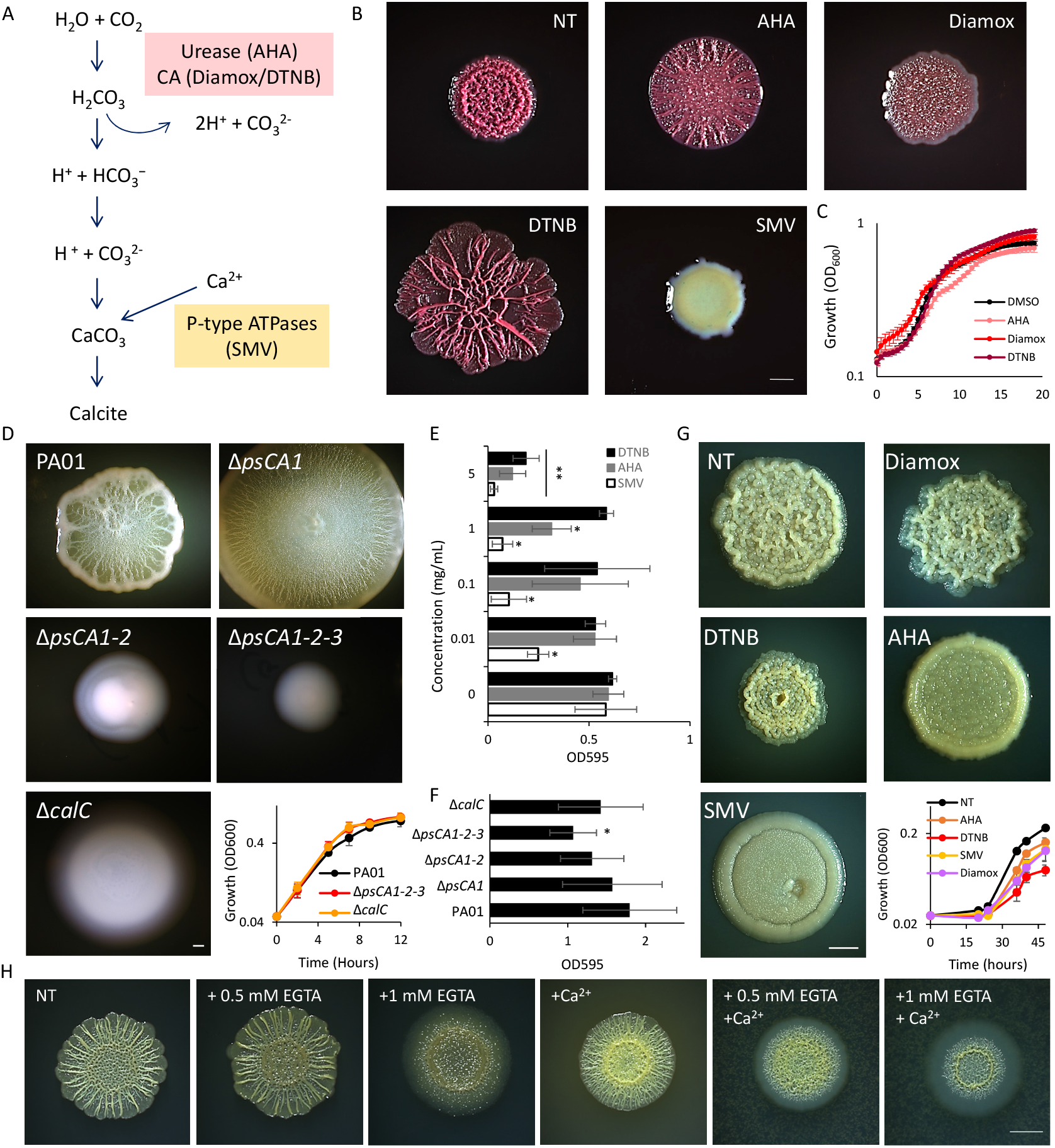
**Inhibition of cellular pathways leading to calcium carbonate production disrupts biofilm development in *P. aeruginosa* and *M. abscessus*.** A. A schematic representation of the chemical reaction leading to calcium carbonate production, with the inhibitors used in this study indicated in parenthesis next to the step they inhibit. B. Light microscopy images of 3-day-old *P. aeruginosa* PA14 biofilm colonies grown on TB agar containing the inhibitors indicated in (A) as follows: untreated (NT), 1.5 mg/ml DTNB, 1.75 mg/ml AHA, 2.5 mg/ml Diamox and 0.01 mg/ml SMV. Scale bar – 2 mm. A representative image (out of n = 3 experiments) is shown. C. The effect of inhibiting calcium carbonate production on planktonic growth of *P. aeruginosa* PA14. Bacteria were grown shaking in liquid TB medium, supplemented with: 1.5 mg/ml DTNB, 1.75 mg/ml AHA, 2.5 mg/ml Diamox and 0.01, 0.05 and 0.01 mg/ml SMV, or untreated (NT). Growth was monitored by measuring OD600 in a microplate reader every 30 min. Results are averages of six wells, bars represent standard deviations. A representative experiment (out of n= 3 experiments) is shown. D. Light microscopy images of 4-day-old biofilm colonies of wild type *P. aeruginosa PA01* and indicated mutants grown on BHI agar: carbonic anhydrases (*psCA1* – PA0102, *psCA2* - PA2053, *psCA3* – PA4676) and calcium channel (Δ*calC*). A representative experiment (out of n= 3 experiments) is shown. Insert – the indicated mutant strains were grown planktonically in shaking liquid BHI medium. Growth was monitored by measuring OD600 in a spectrophotometer at indicated times. Results are averages of 4 wells, bars represent standard deviations. Scale bar – 2 mm. A representative experiment (out of n= 3 experiments) is shown. E. Crystal violet assay quantifying biofilm formation of *P. aeruginosa* PA14, grown in BHI supplemented with the indicated inhibitors. Results are averages and standard deviation of three independent experiments. P values, as determined by student’s t-test are indicated (* pVal <0.01, ** pVal <0.001, when compared to the untreated control). F. Crystal violet assay quantifying biofilm formation of wild type and indicated mutant derivates of *P. aeruginosa* PA01, grown with in liquid BHI medium. Results are averages and standard deviation of three independent experiments. P values, as determined by student’s t-test are indicated (* pVal <0.01, when compared to the untreated control). G. Light microscopy images of 5 day-old *M. abscessus* biofilm colonies, grown on B4 agar supplemented with calcium and 2.5 mM of the indicated inhibitors or untreated (NT). A representative experiment (out of n = 3) experiments is shown. Insert – *M. abscessus* was grown planktonically in liquid B4 medium. Growth was monitored by measuring OD_600_ in a spectrophotometer at indicated times. Scale bar – 2 mm. Results are averages of three independent repeats, bars represent standard deviations. H. Light microscopy images of 3-day-old *P. aeruginosa* PA14 biofilm colonies grown on SCFM agar, either untreated (NT) or suplemented with 700 mM CaCl_2_ and EGTA, as indicated. Scale bar – 5 mm. A representative experiment (out of n = 3) experiments is shown.

The cellular pathways of biomineralization present in all three bacteria are strikingly conserved (Fig. 4A), and can be inhibited at several points. The role of urease and carbonic anhydrase (CA) in biomineralization is well established (Jansson and Northen, 2010; Phoenix and Konhauser, 2008). The consequent action of those enzymes increases local bicarbonate levels, and at the same time increases the pH, creating an environment favorable to mineral precipitation. In the section above, we have shown that intracellular calcium was needed for calcite formation. Consistent with the central role of calcite scaffolds in supporting biofilm morphology, inhibition of urease, CA, and especially P-type ATPases, disrupted *P. aeruginosa* biofilm morphology (Fig. 4B) and abolished the mineral macrostructures as judged by microCT X-Ray (Supplementary Fig. 15). This effect could not be explained by inhibition of planktonic growth, which was mostly unaffected (Fig. 4C, Supplementary Fig. 16). Similarly, reducing self-produced and atmospheric carbon dioxide reduced the mineral macrostructure (Supplementary Fig. 17).

**Figure 5:**
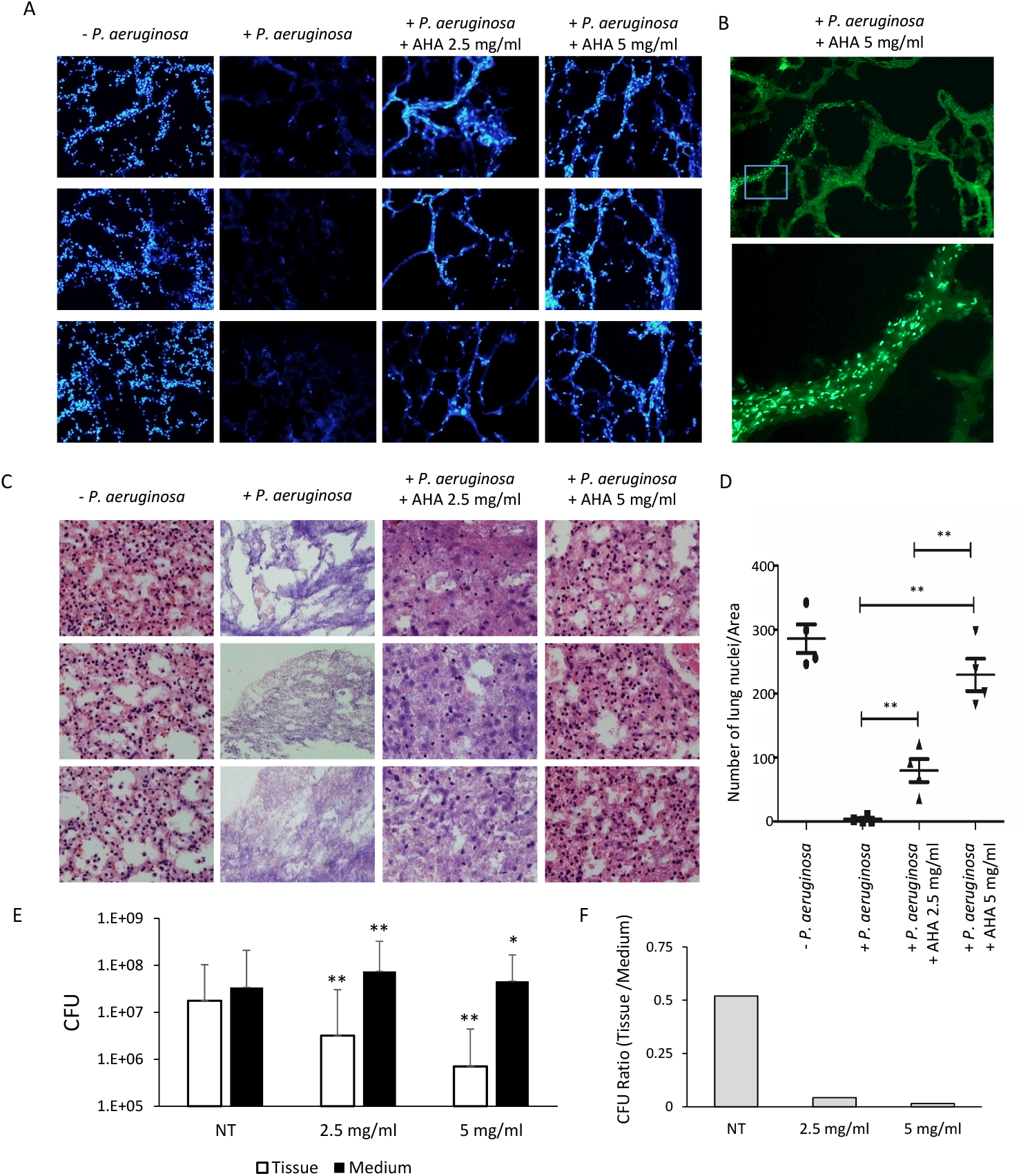
**Inhibition of urease prevents biofilm formation and tissue deterioration in an *ex-vivo* lunge model** A. and B. Fluorescent microscopy images of 10 micron slices of lung tissues, infected with *P. aeruginosa* PA14 and treated with AHA, as indicated. (A) Blue – lung cell nuclei stained by DAPI, (magnification x20). (B) Green - GFP expressing *P. aeruginosa*, (magnification x40). A representative experiment (out of n = 3) experiments is shown. C. Light microscopy images of 7 micron slices of lung tissue. Samples were treated as in (A), and stained with hematoxylin and eosin (H&E). Blue – nuclei, red – extracellular matrix and cytoplasm, (magnification x40). A representative experiment (out of n = 3) experiments is shown. D. Quantification of lung viability. Lung tissue cultures were infected with *P. aeruginosa* PA14 and treated with AHA, as indicated. ImageJ 1.51g software was used to automatically count lung cell nuclei in 4 randomly chosen fields for each treatment. P values, as determined by student’s t-test are indicated (** pVal <0.001). E. CFU analysis of *P. aeruginosa* PA14 infecting lung tissue and surrounding media, with and without treatment. P values, as determined by student’s t-test are indicated (* pVal <0.01, ** pVal <0.001, when compared to the untreated control). F. Quantification representing the ratios of tissue to medium CFU in (E).

**Figure 6:**
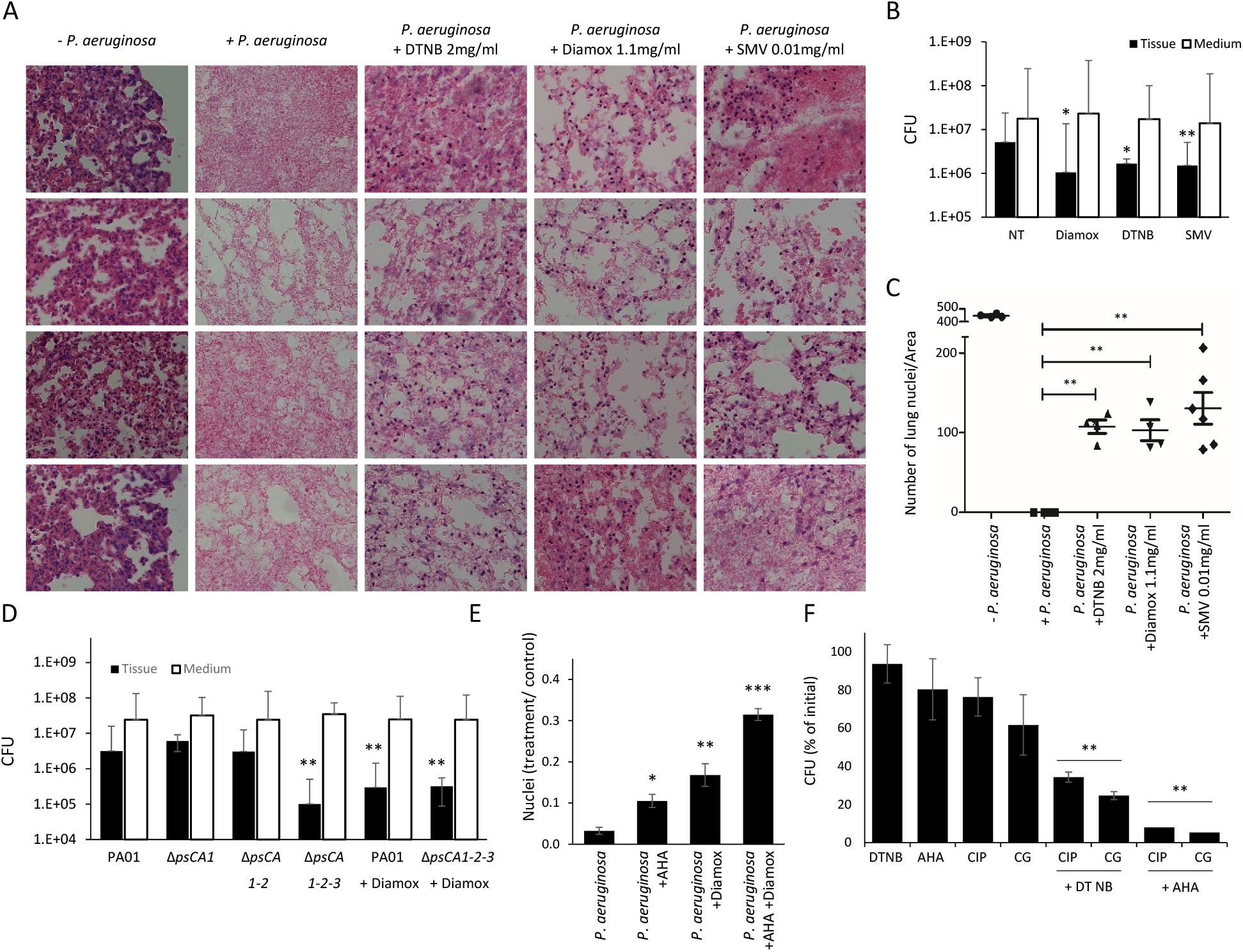
**Inhibition of calcium carbonate production prevents biofilm formation and tissue deterioration in an *ex-vivo* lunge model** A. Light microscopy images of 7 micron slices of lung tissues infected with *P. aeruginosa* PA14 and treated with biomineralization inhibitors as indicated. Samples were stained with H&E. Blue – nuclei, red – extracellular matrix and cytoplasm, (magnification x40). A representative experiment (out of n = 3) experiments is shown. B. CFU of *P. aeruginosa* PA14 infecting lung tissue culture and surrounding media with and without indicated inhibitors (as in A). P values, as determined by student’s t-test are indicated (* pVal <0.01, ** pVal <0.001, when compared to the untreated control). C. Quantification of lung viability. Lung tissue cultures were infected with *P. aeruginosa* PA14 and treated with biomineralization inhibitors as indicated. ImageJ 1.51g software was used to automatically count lung cell nuclei in 4 randomly chosen fields for each treatment. P values, as determined by student’s t-test are indicated (** pVal <0.001). D. CFU analysis of wild type *P. aeruginosa* PA01 and the indicated mutants infecting lung tissue culture and surrounding media. Diamox was applied at 1.1 mg/ml, when indicated. P values, as determined by student’s t-test are indicated (* pVal <0.01, ** pVal <0.001, when compared to the untreated control). E. Quantification of lung viability. Lung tissue cultures were infected with *P. aeruginosa* PA14 and treated with biomineralization inhibitors, as indicated (Diamox and AHA applied at 1.1 mg/m). ImageJ 1.51g software was used to automatically count lung cell nuclei in 4 randomly chosen fields for each treatment. P values, as determined by student’s t-test are indicated (** pVal <0.001). F. *P. aeruginosa* PA14 biofilms were grown for 72 h on TB agar. The colonies were physically divided into equal parts, harvested, and half was treated for 2 h, as indicated. CIP – Ciprofloxacin, 3 mg/ml; GC – gentamycin, 10 mg/ml; AHA – 2.5 mg/ml; DTNB – 2 mg/ml. Viability is expressed as the ratio of CFU between the treated and the untreated parts of a colony. Results are averages and standard deviation of three independent experiments. P values, as determined by student’s t-test are indicated (** pVal <0.001).

Inhibition of complex calcium-dependent colony formation, but not planktonic growth, was also observed in mutants lacking two or more carbonic anhydrase genes or a calcium transporter CalC (Fig. 4D). Biofilm formation on polystyrene, as quantified by crystal violet staining, was also severely compromised by either chemically or genetically impairing carbonate formation or calcium uptake (Fig. 4E-F). The fact that SMV addition or CalC deletion could prevent 3D morphology development and biofilm formation, but had little effect on planktonic growth, is consistent with our view that carbonate mineralization in biofilms relies on regulation of intracellular calcium homeostasis.

Potentially broad clinical implication of these findings was supported by testing the same chemical inhibitors on *M. abscessus*. Colony morphology could be compromised by inhibiting biomineralization enzymes (Fig. 4G and Supplementary Fig. 19). Morphology defects could not be explained by inhibition of cell growth, as most potent morphology inhibitors AHA and SMV had the smallest effect on planktonic growth (Fig. 4G). We could therefore conclude that biomineralization is indeed highly conserved, and is mediated by a previously untargeted druggable pathway.

Biofilm mineralization was not previously demonstrated in the lung microenvironment. To test the ability of lung pathogens to form structured communities in such conditions, we grew *M. abscessus* and *P. aeruginosa* on synthetic CF sputum medium (SCFM). Both species were capable to form highly organized 3D colonies on SCFM (Supplementary Fig. 20). For *P. aeruginosa*, this robust colony architecture was enhanced by addition of calcium to the growth medium, and diminished with the addition of calcium chelator EGTA (Fig. 4H); suggesting that biofilms may rely on crystalline calcium carbonate scaffolds to support the community architecture in the lung. Therefore, we examined whether calcite formation by lung pathogens could be also observed in clinical samples. We collected sputum samples from CF patients chronically infected with *P. aeruginosa* and removed the organic matter by bleach. Insoluble material indicating the presence of a mineral was found in 31 out of 50 samples examined (Supplementary Table 3). These putatively positive patients were further examined by FTIR for residual pure minerals, and spectra typical of calcite, lacking a ν1 peak and containing a ν_2_ peak at 875 cm^-1^, ν_3_ peak at 1425 cm^-1^, and ν_4_ peak at 713 cm^-1^ (Supplementary Figure 21), were identified in 7 samples.

To further explore the role of biomineralization during lung infection by *P. aeruginosa*, we set up a lung *ex vivo* model. Organ cultures derived from lung and colon tissues have been applied previously for the study of biofilm infections and viral tropism (Kolodkin-Gal et al., 2008; Kolodkin-Gal et al., 2007; Massler et al., 2011; Wu et al., 2021), and thus can potentially be used to study lung infections. We cultured mice lungs and then infected them with *P. aeruginosa* constitutively expressing GFP. Fluorescent microscopy revealed biofilms forming on the tissue prior to its destruction (Supplementary Fig. 22). Using this system, we then tested the effect of chemically blocking biomineralization during initial infection stages. When left untreated, *P. aeruginosa* infected tissue (labeled by DAPI for visualization) was completely deteriorated. Infection in the presence of urease inhibitor AHA prevented biofilm formation and rescued the lung tissue (Fig. 5A). While the tissue was rescued, the bacterial load in the growth media remained similar (Fig. 5B). This is consistent with the *in vitro* results presented above – as in all cases inhibition of biomineralization prevented biofilm formation, but had little effect on planktonic growth. This further strengthens our proposed hypothesis, suggesting biomineralization is a specific and controlled developmental feature of bacterial biofilms. To quantify the effects of urease inhibition during lung tissue infection, the tissue was stained with H&A and the nuclei were counted (Fig. 5C-D). We observed a significant rescue of the infected tissue by AHA, in a dose-dependent manner. In agreement with these findings, the growth of bacteria in the medium was not reduced by AHA addition, but their colonization of the lung tissue was largely compromised (Fig. 5E and 5F).

Significant reduction of lung cell death during infection by *P. aeruginosa* was also obtained with CA and calcium uptake inhibitors (Fig. 6A-C), and with mutants lacking CAs (Fig. 6D). Consistent with our observation that both Diamox and AHA inhibited lung colonization and subsequent cell death, the combination of both was additive (Figure 6E). At the concentrations used here, AHA and DTNB were slightly toxic to the uninfected lung tissue, while SMV and Diamox had no adverse effect (Supplementary Fig. 23). We can therefore conclude that the entire biomineralization pathway, rather than a specific enzyme, is of crucial importance to host infection.

Finally, while the ability to rescue host tissue by inhibition of biomineralization is encouraging, in clinical settings the eradication of bacteria is essential. We therefore tested whether inhibition of mineralization during lung infection of *P. aeruginosa* could increase its sensitivity to antibiotics. While neither biomineralization inhibitors nor quinolone antibiotics (a first-line antibiotic for treating *P. aeruginosa* infections) (Hewer and Smyth, 2017) could effectively eradicate established biofilms, the combination of both classes of drugs significantly increased the sensitivity of biofilms to treatment (Fig. 6F) – highlighting the potential clinical applications of biomineralization inhibition.

## Discussion

Bacterial biofilms are widespread in nature, forming multicellular colonies on various surfaces. The molecular mechanisms guiding the complex network of events leading to the transition from a free-living planktonic bacterium to the differentiated community are well characterized. Much is known about the production of organic matrix, the division of labor between the different cell types, the metabolic adaptations, the methods of communication between the community members – and the fitness advantages of such a communal life style are evident.

In addition, bacterial biofilms were extensively shown to promote biomineralization of metals (such as gold and iron), phosphate and carbonate (Phoenix and Konhauser, 2008). In this work, we uncover a previously overlooked, but fundamental, connection between the highly regulated biofilm development and the seemingly random biomineralization occurring within biofilms. We propose that bacterial communities actively control the formation of mineral macrostructures and that those “skeletons” play an important role in biofilm development and fitness.

Moreover, the conserved and wide-spread nature of biomineralization offers novel therapeutic approaches to target one of the most challenging global threats to human health – drug-resistant chronic biofilm infections.

The first clues to the presence of a structural mineral component in a biofilm came when we demonstrated that a mature 3D structure of biofilm colonies of *B. subtilis* depends on their ability to precipitate precisely organized patterns of calcium carbonate (Oppenheimer-Shaanan et al., 2016). A controlled deposition of a calcium carbonate scaffold structurally supports the complex architecture of biofilm colonies and protects them from the environment by limiting solute diffusion (Keren-Paz et al., 2018).

In this work, we build on those preliminary observations to show that crystalline calcium carbonate production is a controlled process fundamental to biofilm development. Only when calcification occurs, *B. subtilis* can successfully activate and maintain the characteristic biofilm transcriptional program, activating matrix production and sporulation while repressing motility (Fig. 1).

Mechanistically, the formation of mineral macro-structures is initiated within intracellular compartments of a small subpopulation of cells, which secrete small calcium-rich granules, while remaining intact (Fig. 2). The process is likely actively controlled, as the membrane integrity of the exporting cells is not compromised. Initiating mineral formation within the controlled cellular environment allows the organism to regulate the composition, structure, and physical properties of the mineral – therefore controlling the final function.

The best known example of bacteria producing functional minerals formed in a defined intracellular compartment by a highly regulated cellular pathway is the magnetotactic bacteria, forming invagination of membranes to generate magnetosomes (Komeili et al., 2006). Recently, intracellular calcium carbonate mineralization was reported in cyanobacteria (Blondeau et al., 2018) and in a subgroup of poorly characterized magnetotactic bacteria (Monteil et al., 2020). Unlike the above examples, which occur in a specific group of phylogenetically related planktonic aquatic bacteria, the formation of mineralized macrostructures in biofilms is conserved across the microbial domain, highlighting their potential importance for general microbial fitness, unrelated to specific adaptations of single-cell microbial metabolism.

Regulated calcification was found to mediate complex 3D biofilm structure in three phyla – the Gram-positive spore former *B. subtilis* and actinobacterium *M. abscessus*, and the Gram-negative *P. aeruginosa* (Fig. 3). The conserved nature of cellular pathways promoting biomineralization allowed us to prevent biomineralization in phylogenetically distinct bacteria by chemically and genetically inhibiting carbonate accumulation and calcium uptake (Fig. 4). These results support the notion that mineralization in biofilms is a well-defined developmental process, initiated by a dramatic change in the structure and function at the single cell level within a subpopulation of cells in bacterial biofilms. Similar to the formation of bones initiated by the osteoblast cells within higher animals, the presence of mineral-producing cells within a biofilm leads to the formation of a macrostructure serving specific functions and increasing the fitness of the microbial community.

In this work, we demonstrate that, in a subpopulation of cells, calcium uptake and functionally conserved pathways of carbonate production alter the intracellular microenvironment, promoting mineral nucleation and leading to formation of a mineral macro-structure. Numerous studies of *B. subtilis, P. aeruginosa* and additional biofilm formers, established that complex colony architecture is dependent on the organic extracellular matrix (Branda et al., 2005; Colvin et al., 2012; Jones and Wozniak, 2017; Kobayashi and Iwano, 2012; Romero et al., 2010; Vlamakis et al., 2013), and thus hypothesized that the presence of the organic ECM components is sufficient for bacterial biofilm morphogenesis. We found that non-organic ECM components are also necessary for the development of structured bacterial communities. While our results indicate that mineral formation initiates within the cells and requires a controlled intracellular environment, the final mineral macro-structure assembly has to occur in the extracellular microenvironment, guided by complex interactions with the extracellular organic matrix (Keren-Paz and Kolodkin-Gal, 2020), as indicated by the scanning electron microscopy (Figure 4C, (Oppenheimer-Shaanan et al., 2016)). Therefore, both organic ECM components and biogenic minerals are essential but not sufficient for the formation of complex structures.

While in our settings bacteria relied on calcite to generate functional scaffolds, other minerals may serve similar functions. For example, deposition of calcium phosphate crystals in *P. mirabilis, P. vulgaris* and *Providencia rettgeri* biofilms blocks catheters in infected patients (Broomfield et al., 2009; Mathur et al., 2006; Tan et al., 2018), and urease-dependent calcium phosphate mineralization was shown to increase *P. mirabilis* resistance to ciprofloxacin (Li et al., 2016c) and increase its survival in dual-species biofilms (Li et al., 2016b). Importantly, the physiological role of calcium phosphate deposits within these medical biofilms and whether dedicated cells facilitate this process remains to be determined.

Intriguingly, we could observe the presence of bacterially produced calcite minerals in the sputum of some CF patients infected with *P. aeruginosa*. This observation raises the possibility that lung pathogens can generate functional minerals within the host. While biofilm colonies formed on top of biofilm-inducing agar are a convenient model system mimicking many of the environmental and physical conditions of the host, it lacks the host itself. While several murine models are available for the study of CF (such as *Cftr-/-* knockout mice), they all fail to replicate the acute lung disease and chronic bacterial infections characteristic of CF patients (Semaniakou et al., 2018). *Ex vivo* organ cultures are an experimental system that recapitulates the three-dimensional structure of the tissue and includes several cell types and a variety of extracellular matrix components. On the other hand, organ culture is a controlled microenvironment lacking compounding factors that play a role in any infection, such as efficiency of drug delivery and the immune system (Kolodkin-Gal et al., 2007).

Using chemical inhibitors and mutants, we demonstrated that biomineralization by *P. aeruginosa* was necessary for lung deterioration, and that preventing it could rescue the infected tissue. The fact the free-living bacteria do not seem to be affected by inhibiting biomineralization, and remain in the culture, while the tissue itself is rescued, highlights the importance of host attachment during infection. It is also possible that the restrictive properties of crystalline minerals on the diffusivity and viscoelasticity could be deleterious to the host tissue. Moreover, consistent with the protective role of the mineral for the bacterial biofilm, chemical or genetic inhibition of mineralization restored the sensitivity the *P. aeruginosa* to antibiotic treatment (Fig. 5 and Fig. 6).

Our work suggests that it is time to reconsider the old definition of bacterial mineralization solely as a passive, unintended byproduct of environmental bacterial activity. Instead, here we provide evidence that it can also be a regulated developmental process originating from a dedicated, previously uncharacterized microbial compartment, conferring clear benefits to biofilm-residing bacteria. Biomineralization is highly conserved, and thus is of enormous clinical significance, as it can yield completely novel classes of broad-spectrum drugs to combat emerging biofilm infections.

## Methods

### Strains and media

All strains used in this work were either wild type or derivatives from *Bacillus subtilis* NCIB 3610, *Pseudomonas aeruginosa* PA14, PA01 and *Mycobacterium abscessus* ATCC 19977 (recently renamed *Mycobacteroides abscessus*). A list of strains used in this study can be found in Supplementary Table 1. Deletions of *arsR*, *mntR*, *zur*, *cueR*, *copA* and *chaA* were generated by transforming *B. subtilis* NCIB 3610 with genomic DNA isolated from *B. subtilis* 168 deletion library {Koo, 2017 #6} (Addgene, Cat#1000000115), and verified by PCR. Deletion of *yetJ* and *yloB* was done as previously described, and double and triple mutants were generated by transformation with genomic DNA, and verified by PCR. *P. aureginosa* PA01 single *psCA1* (CA1), double *psCA1-psCA2* (CA1-2), and triple *psCA1-psCA2-psCA3* (CA1-2-3) mutants were reported earlier (Lotlikar et al., 2019). Deletion of *calC* was generated as described in (Hoang et al., 2000).

### Biofilm assays

*B. subtilis* and *M. abscessus* biofilms were grown on B4 biofilm-promoting solid medium (0.4% yeast extract, 0.5% glucose, and 1.5% agar) (Barabesi et al., 2007) supplemented with calcium acetate at 0.25% v/v, incubated at 30°C and 37°C respectively, in a sealed box for enriched CO_2_ environment achieved by using the candle jar method (Oppenheimer-Shaanan et al., 2016). *P. aeruginosa* biofilms were grown on TB medium, as described previously (Dietrich et al., 2013) or on Brain Heart Infusion Broth (BHI, Sigma-Aldrich) supplemented with 1% agar. SCFM medium was prepared as in (Palmer et al., 2005) and supplemented with 1.5% agar. Crystal-violet biofilm assay was carried out as previously described (O’Toole, 2011).

When indicated, the medium was supplemented with the following inhibitors purchased from Sigma-Aldrich: acetohydroxamic acid (AHA) (Cat. #159034); 5,5’-Dithiobis(2-nitrobenzoic acid) (DTNB) (Cat. #D8130); acetazolamide (Diamox) (Cat. #A6011) and sodium metavanadate (SMV) (Cat. #590088). The concentrations used are indicated in the text.

### Phase microscopy

Biofilm colonies were observed using a Nikon D3 camera or a Stereo Discovery V20" microscope (Tochigi, Japan) with objectives Plan Apo S × 0.5 FWD 134 mm or Apo S × 1.0 FWD 60 mm (Zeiss, Goettingen, Germany) attached to Axiocam camera, as required. Data were captured using Axiovision suite software (Zeiss).

### Planktonic growth assays

All strains were grown from a single colony isolated over lysogeny broth (LB) plates to a mid-logarithmic phase of growth (4 h at 37°C with shaking). Cells were diluted 1:100 in 150 μl liquid B4 medium in 96-well microplate (Thermo Scientific). Cells were grown at 30°C for 20 h in a microplate reader (Synergy 2, BioTek), and the optical density at 600 nm (OD_600_) was measured every 15 min. Three independent experiments were conducted, with three technical repeats per plate.

CFU assay was carried out as previously described in detail (Bucher et al., 2016).

### RNA extraction and library preparation

Biofilm colonies were grown on biofilm-promoting B4 solid medium with and without calcium for 1, 2, 3 and 6 days. Three independent experiments were conducted, with three colonies from each treatment combined for RNA extraction in each experiment. The samples were frozen in liquid nitrogen and stored until extraction. Frozen bacterial pellets were lysed using the Fastprep homogenizer (MP Biomedicals) and RNA was extracted with the FastRNA PROT blue kit (MP Biomedicals, 116025050) according to the manufacturer’s instructions. RNA levels and integrity were determined by Qubit RNA BR Assay Kit (Life Technologies, Q10210) and TapeStation, respectively. All RNA samples were treated with TURBO DNase (Life Technologies, AM2238).

A total of 5 ug RNA from each sample was subjected to rRNA depletion using the Illumina Ribo-Zero rRNA Removal Kit (Bacteria, MRZB12424), according to the manufacturers’ protocols. RNA quantity and quality post-depletion was assessed as above. RNA-seq libraries were contracted with NEBNext® Ultra™ Directional RNA Library Prep Kit (NEB, E7420) according to the manufacturer’s instructions. Libraries concentrations and sizes were evaluated as above, and were sequenced as multiplex indexes in one lane using the Illumina HighSeq2500 platform.

### RNAseq processing

Reads were trimmed from their adapter with cutadapt and aligned to the *B. subtilis* genome (subsp. *subtilis* str. NCIB 3610, NZ_CM000488.1) with Bowtie2 version 2.3.4.1 (Langmead and Salzberg, 2012). The number of uniquely mapped reads per gene was calculated with HT-seq (Anders et al., 2015). Normalization and testing for differential expression was performed with DESeq2 v1.16. A gene was considered to be differentially expressed using the following criteria: normalized mean read count ≥ 30, fold change ≥ 3, and adjusted p value < 0.05. First, we tested for differential expression between samples grown with and without calcium separately for each time point; however, since the results for days 1, 2 and 3 were very similar, we joined days 1-3. The crude read count, normalized read count, and the result of the differentially expression tests are available in Supplementary File 1.

### Comparison between growth conditions

We compared our RNAseq expression data to publically available transcriptomes representing 269 different growth conditions (Nicolas et al., 2012). Because that study used microarray platform and not RNAseq, the comparison was performed using the top 10% genes with the highest expression level of every condition and every replicate (383 genes per sample). We than used Jaccard index to measure the overlap between the conditions of the two platforms (i.e. the current study and (Nicolas et al., 2012)). Prior to the analysis, we removed 152 genes that appear among the top 10% in more than 80% of the conditions.

### Scanning electron microscopy and EDX

Biofilm colonies were grown for 1, 3, 6, 10 and 15 days at 30°C on biofilm-promoting B4 solid medium, with or without calcium. The colonies were fixed overnight at 4°C with 2% glutaraldehyde, 3% paraformaldehyde, 0.1 M sodium cacodylate (pH 7.4) and 5 mM CaCl_2_, dehydrated and dried as in (Bucher et al., 2016; Bucher et al., 2015b). Clinical samples were first partially or completely bleached to remove the organic material by 1 hour of incubation in either 3% or 6% sodium hypochlorite, respectively. The insoluble material was collected, washed three times in PBS, three times in acetone and air-dried for 16 hours.

Mounted samples were coated with 15 nm thick carbon layer in carbon coater (EDWARDS). The imaging by secondary electron (SE) or back scattered electron (BSE) detectors and the Energy Dispersive X-ray Spectroscopy (EDS, Bruker) were performed using Carl Zeiss Ultra 55 or Supra scanning electron microscopes.

### Cryo-STEM analysis

Bacterial colonies grown as described were suspended in PBS buffer. Quantifoil TEM grids were glow-discharged with an Evactron Combi-Clean glow-discharge device, and 5 microliters of suspended cells were deposited onto the glow-discharged grids. Ten nm-diameter gold fiducials (Duchesne et al., 2008) were applied before blotting and verification using a Leica EM-GP automated plunging device (Leica). No chemical fixation was used to ovoid artifacts

Vitrified samples were observed with a Tecnai F20 S/TEM instrument (Thermo Fisher Scientific) at 200 kV, with Gatan 805 brightfield and Fischione HAADF detectors. Microscope conditions: extraction voltage = 4300 V, gun lens = 3 or 6, and spot size = 5 or 6 with 10 micrometer condenser apertures, yielding probe diameters of 1-2 nm and semi-convergence angles of ∼ 1.3-2.7 mrad. Images of 2048 X 2048 pixels were recorded with probe dwell times of 8-18 microseconds. Spatial sampling was set between 1 and 4 nm/pixel. Electron doses were 1–3 electrons/A^2^ per image. Single-axis tilt series were recorded using SerialEM (Kremer et al., 1996). EDX was performed in STEM mode on vitrified cell samples with the same electron microscope set-up as used for STEM imaging, using a liquid N_2_ cooled Si(Li) detector (EDAX). Vitrified grids contacting 100-200 cells per grid were automatically imaged on a Talos Arctica (Thermo Fisher Scientific) microscope in STEM mode, and mapped at intermediate magnifications (10-13k) using SerialEM software to obtain large fields of view for quantification of data presented in Fig. 2C. The quantification was done by manually counting the cells with or without dense deposits.

### Tomography reconstructions and visualization

The CSTET tomographic tilt series were aligned using fiducial markers and reconstructed using weighted back projection (Frangakis and Hegerl, 2001) as implemented in the IMOD software suite (Kremer et al., 1996). Reconstructions are displayed after non-linear anisotropic diffusion filtering within IMOD. Segmentation and volume rendering were performed using Amira 6.3 (FEI Visualization Sciences Group).

### STED image acquisition

Cells were isolated from 3-6 day old biofilm colonies and stained with Calcein-AM (20 µM) and NileRed (100 µg/ml). Immediately after the staining procedure, the cells were mounted on a coverslip and immobilized by an agarose pad (1% agarose, S7_50_ minimal media). STED microscopy was performed using a Leica SP8 confocal microscope with a 100x oil immersion objective (NA: 1.4). A 552 nm laser line was used for confocal detection of NileRed, a 494 nm excitation laser and a 592 nm depletion laser were used for G-STED detection of Calcein-AM. Channels were acquired with 200 Hz (4 x line averaging) and the appropriately set HyD hybrid detectors. Images processing was performed using Leica LAS X and the deconvolution of the respective channels was performed using the Huygens algorithm.

### Micro-CT X-ray analysis

Images of indicated magnification were taken using a Zeiss micro XCT 400 instrument (Pleasanton, CA, USA). Tomography was carried out using a micro-focused source set at 20 kV and 100 μA. 1200 separate 2D images were taken with a pixel size of 0.87 mm over 1800, exposure time of 30 sec. Image analysis was carried out with Avizo software (VSG, Hillsboro, OR, USA).

### FTIR spectrophotometer analysis

Calcite was collected as described by Mahamid *et al*., with some modifications (Mahamid et al., 2008): agar samples were slightly bleached with 6% sodium hypochlorite for 1 min to remove organic matter, washed with Milli-Q water twice and dehydrated in acetone.

FTIR spectra of the bleached samples were acquired in KBr pellets by using a NICOLET iS5 spectrometer (Thermo Scientific, Pittsburgh, PA, USA). The samples were homogenized in an agate mortar and pestle with about 40 mg of KBr, and pressed into a 7 mm pellet using a manual hydraulic press (Specac, Orpington, UK). Infrared spectra were obtained at 4 cm−1 resolution for 32 scans.

The infrared calcite spectrum has three characteristic peaks, designated v_2_, v_3_, and v_4._ Calcium carbonate ν_3_ peak is expected at 1425 cm^-1^ for calcite, 1490 cm^-1^ for vaterite and 1475 cm-1 for aragonite. The calcium carbonate ν_2_ peak is expected at 875 cm^-1^ for calcite, 850 cm^-1^ for vaterite and 855 cm^-1^ for aragonite. Finally, the calcium carbonate ν_4_ peak is expected at 713 cm^-1^ for calcite, 750 cm^-1^ for vaterite and 715 cm^-1^ for aragonite (Politi et al., 2004).

### Patient’s samples

Sputum was collected from adult patients as published previously (Schiller and Millard, 1983) and stored in 4°C degrees. All patients positive for calcite were carrying chronic *pseudomonas* infections. Specific patient information can be found in Supplementary Table 2. The samples were collected under Helskinki approval to Prof. Eitan Keren 0456-17-HMO. For FTIR analysis, organic matter was removed as described above.

### *Ex vivo* lung infection system

Animal work was carried out under ethical approval MD-16-15035-1. Lungs were harvested from 2 mice (one month old) and placed in petri dishes containing DMEM 5% FCS. We divided the tissue into circular pieces 3 mm in diameter with a biopsy punch and transferred to a 24 well plate (4-5 explants/well) with 450 µl DMEM. DMEM was applied with carbenicillin 100 µg/ml (Sigma-Aldrich) and chemical inhibitors, as indicated. To each respective well, either 50 µl DMEM (control) or *P. aeruginosa* (pretreated with chemical inhibitors when indicated and diluted within DMEM to attain OD_600_ 0.4) was added – with three technical repeats for each condition. The plates were incubated at 37°C for ∼2 days (52 hours), washed twice with PBS, fixed with PFA 4% for 10 minutes, and imbedded in either cryosection (OCT) compound or paraffin. Paraffin samples were cut into 7 micron slices and stained with H&E; whereas cryosections were cut into 10 micron slices and placed on superfrost plus slides.

### Cultivable Cell Quantification (CFU)

CFU quantification was done as previously described (Bucher et al., 2015a). For CFU of bacteria infecting *ex vivo* tissue culture: individual punches or their growth media were collected, Punches were resuspended in 1 ml PBS (Biological Industries), and free-living bacteria were pelleted and resuspended in PBS at the same volume. Samples were thoroughly vortexed (5 min). The samples were then mildly sonicated (BRANSON digital sonifier, Model 250, Microtip, amplitude 30%, pulse 2 × 5 s). In all cases, to determine the number of viable cells, samples were serially diluted in PBS, plated on LB plates, and colonies were counted after incubation at 30°C overnight. For

## Acknowledgments

The Kolodkin-Gal lab is supported by the Israel Science Foundation grant number 119/16, Israel Foundation grant number JSPS 184.20, Kamin grant by Israel Chief Scientist no. 67459, Israel Ministry of Science, Technology & Space (grant no. 713454), Ministry of Health (grant no. 713645), Angel-Fiavovich fund for ecological research, Dr. Barry Sherman Institute for Medicinal Chemistry, Kekst Family Institute for Medical Genetics and by the Helen and Milton A. Kimmelman Center for Biomolecular Structure & Assembly. IKG is a recipient of Rowland and Sylvia Career Development Chair. The electron microscopy studies were partially supported by the Irving and Cherna Moskowitz Center for Nano and BioNano Imaging (Weizmann Institute of Science). We thank Prof. Ehud Banin and to Itzhak Zander (Bar-Ilan University) for helpful discussions, and for providing PA01 strains.

## Supplementary figures

**Supplementary Figure 1.**
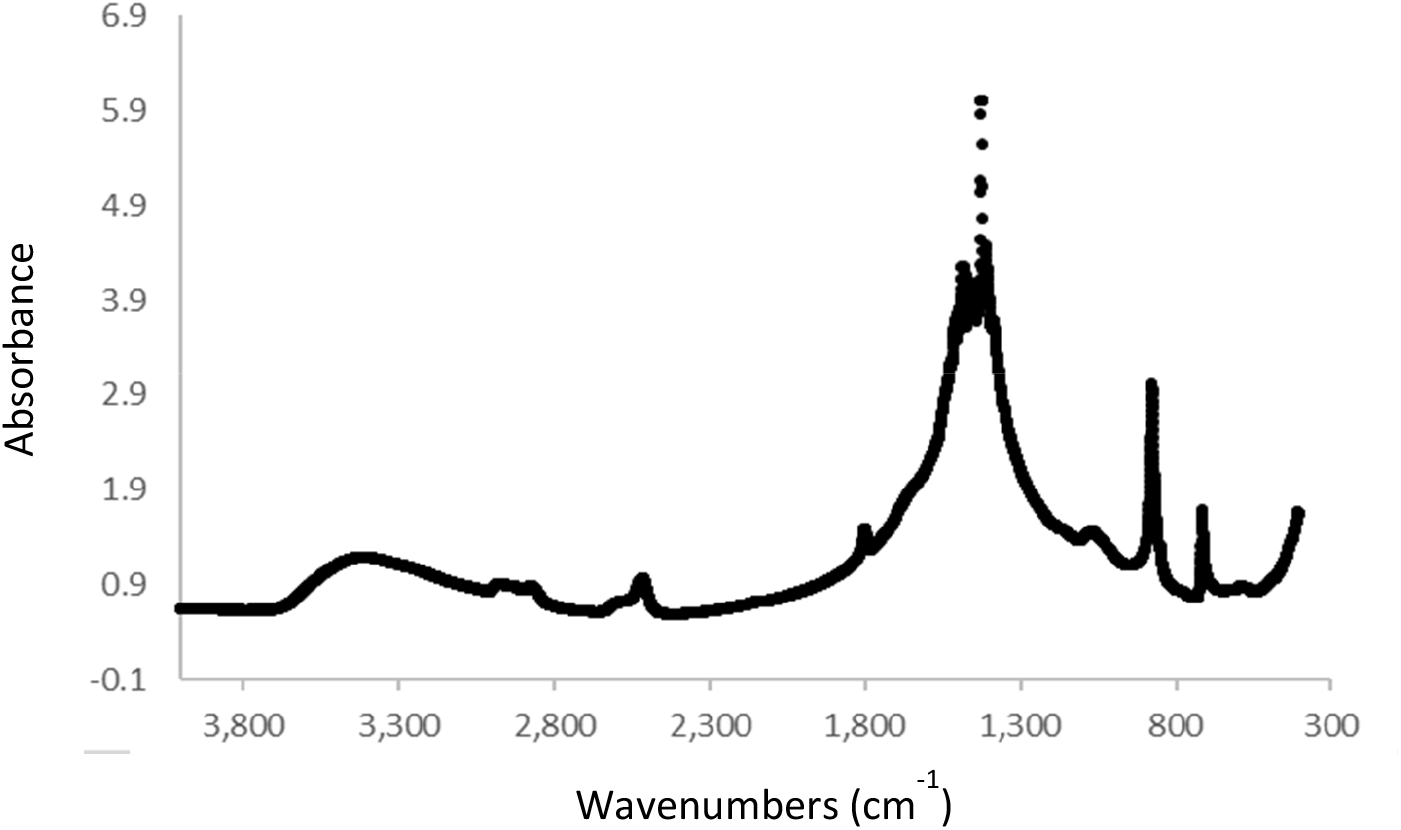
FTIR spectra of bleached *B. subtilis* biofilm colonies, displaying vibrations characteristic of calcite (crystalline calcium carbonate).

**Supplementary Figure 2.**
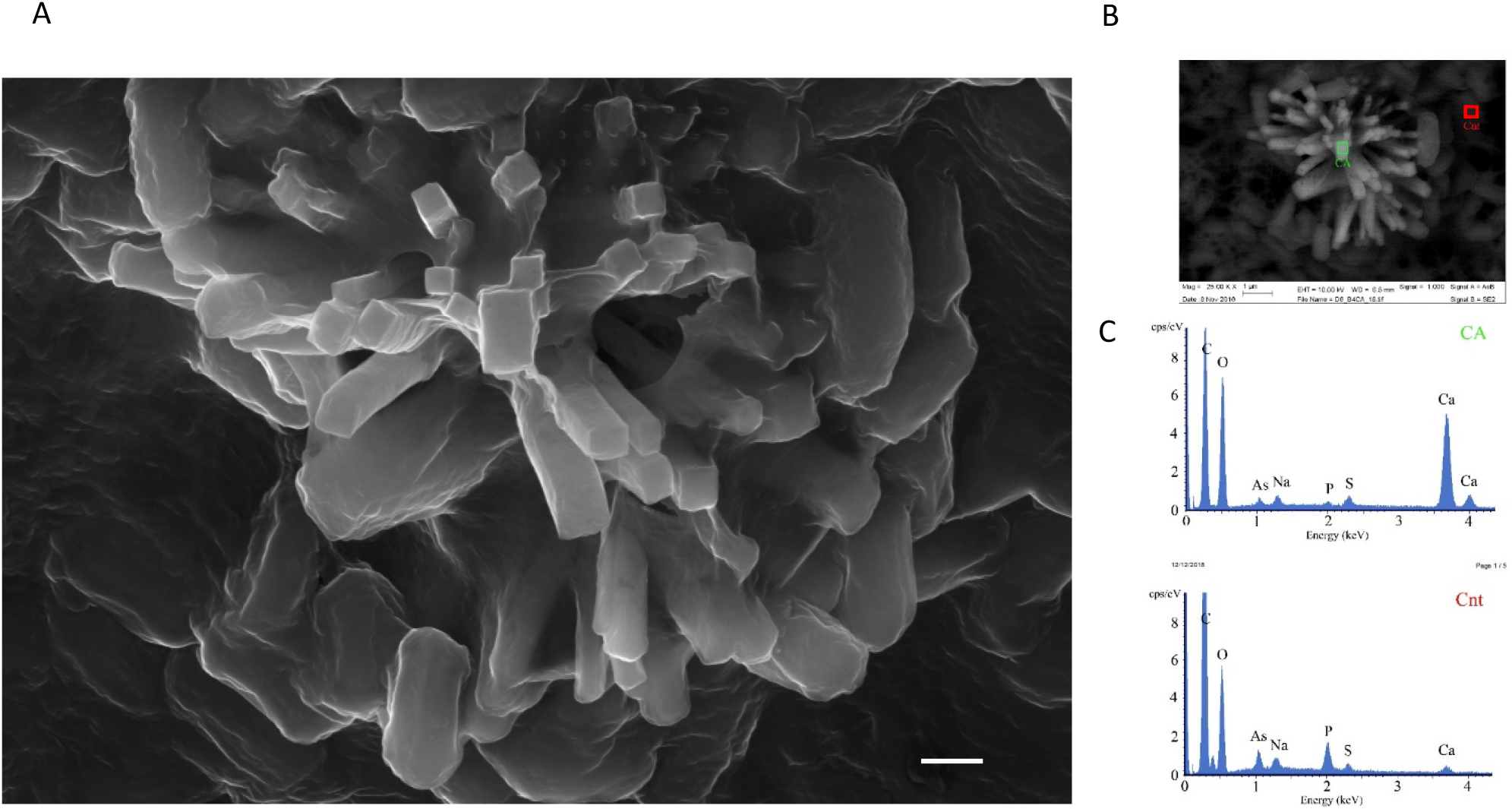
(**A**). Scanning Electron Microscopy (SEM) image of *B. subtilis* biofilm colony grown for 6 days on a biofilm-inducing B4-Ca^2+^ agar. Shown are the surface of the biofilm and rhombohedral mineral structures in secondary electron mode. Magnification – X25000, scale bar – 1 µm. **(B).** Backscattered mode SEM image of the area shown in (**A**). **(C)** EDX spectra of the area shown in (**A**). CA – calcium rich area; Cnt – control area. A representative field (out of n=4 fields, from 8 independent experiments) is shown.

**Supplementary Figure 3.**
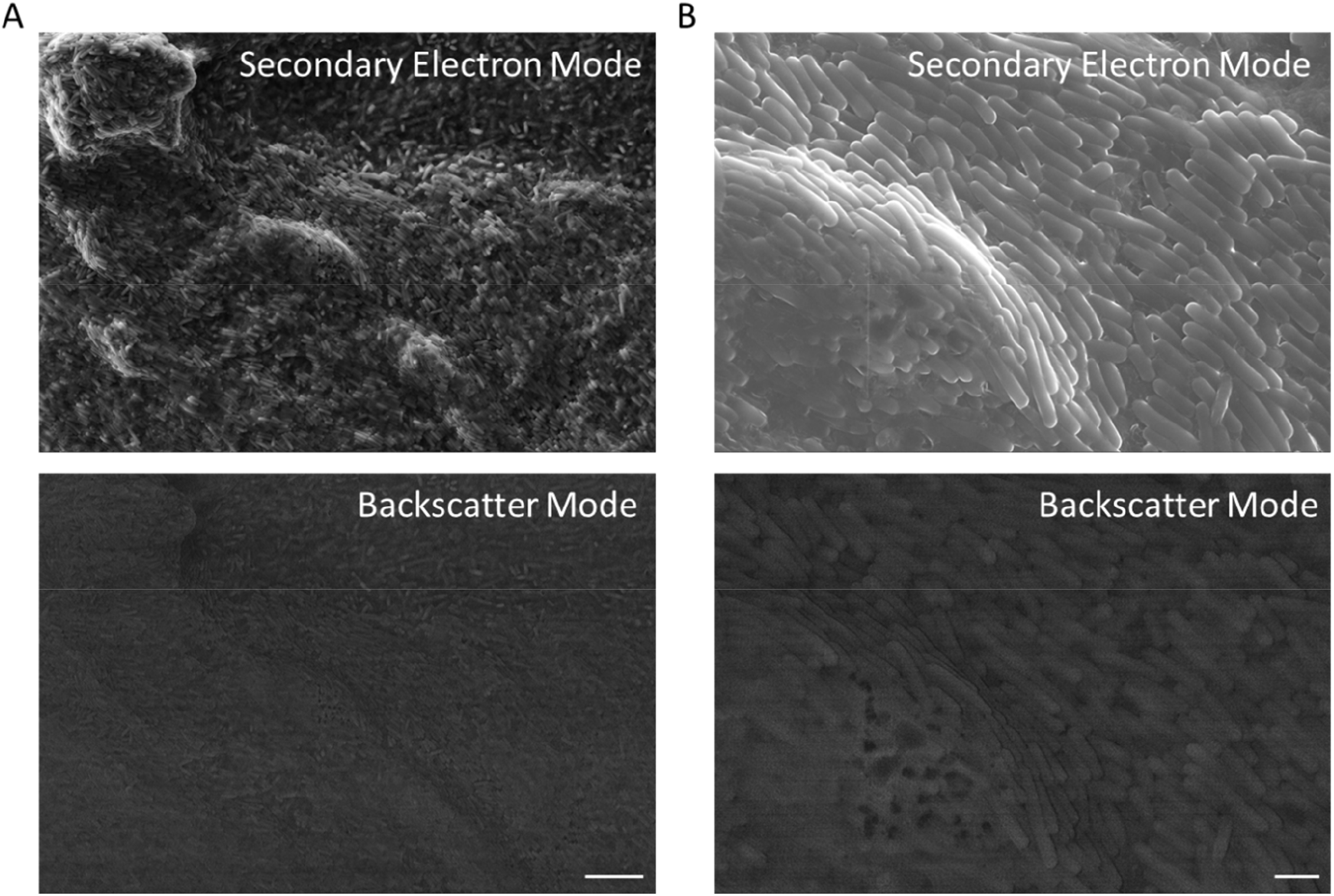
Scanning Electron Microscopy (SEM) image of *B. subtilis* biofilm colony grown for 6 days on a B4 agar (without excess calcium). Shown are the surface of the biofilm in secondary electron and in backscattered mode of a representative field. **(A)** Magnification – X2500, scale bar – 10 µm. **(B).** Magnification – X10000, scale bar – 2 µm. Representative fields (out of n=4 fields, from 8 independent experiments) are shown.

**Supplementary Figure 4.**
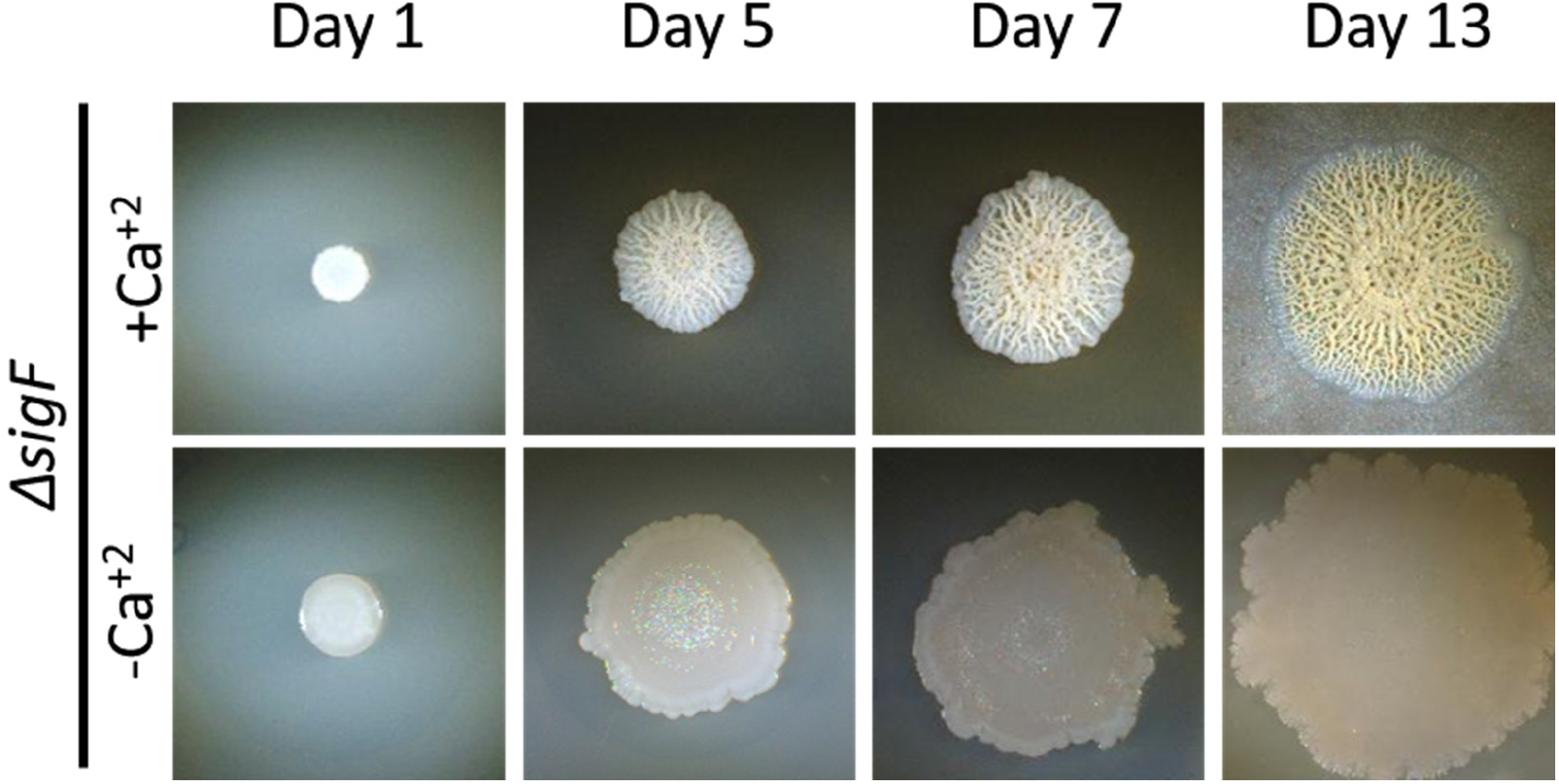
Light microscopy images of Δ*sigF* mutant strain. Biofilm colonies were grown on B4-Ca^2+^ agar for the indicated time at 30°C. The experiment was repeated 3 times, in a technical quadruplicate – and representative images are shown.

**Supplementary Figure 5.**
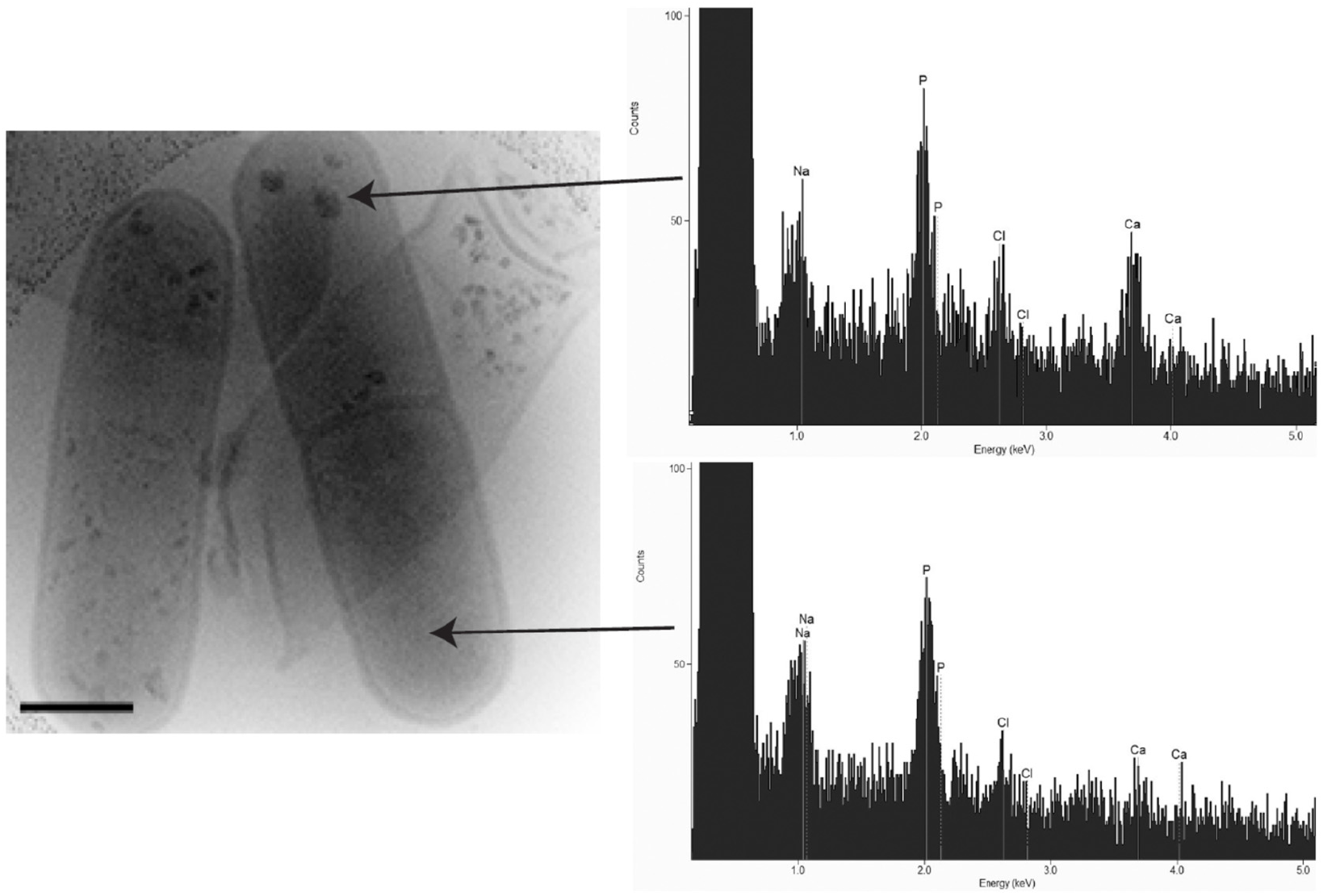
Left panel - Bright-field STEM image of representative *B. subtilis* biofilm cells from a showing cellular calcium deposits, from a colony grown 10 days on B4-Ca^2+^ agar. The black arrow indicates the cell used for EDX analysis in panels (B-C). Right panel – EDX analysis of the calcium deposits. A representative mineralizing cell imaged (out of n=50 cells, from 4 independent experiments performed with EDX is shown.

**Supplementary Figure 6.**
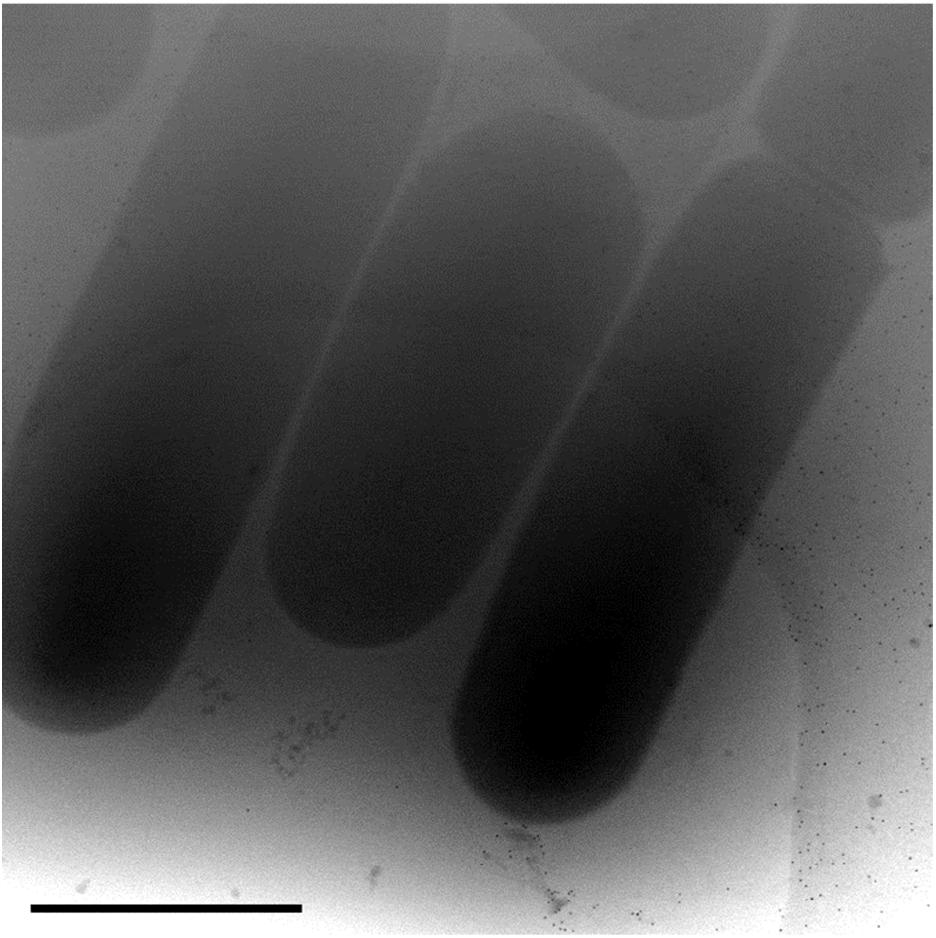
Cryo-STEM bright-field images of representative biofilm cells from a *B. subtilis* colony grown for 10 days on B4 agar (without calcium). Scale bar 1 µm. A representative field imaged (out of n=30 cells, from 4 independent experiments is shown.

**Supplementary Figure 7.**
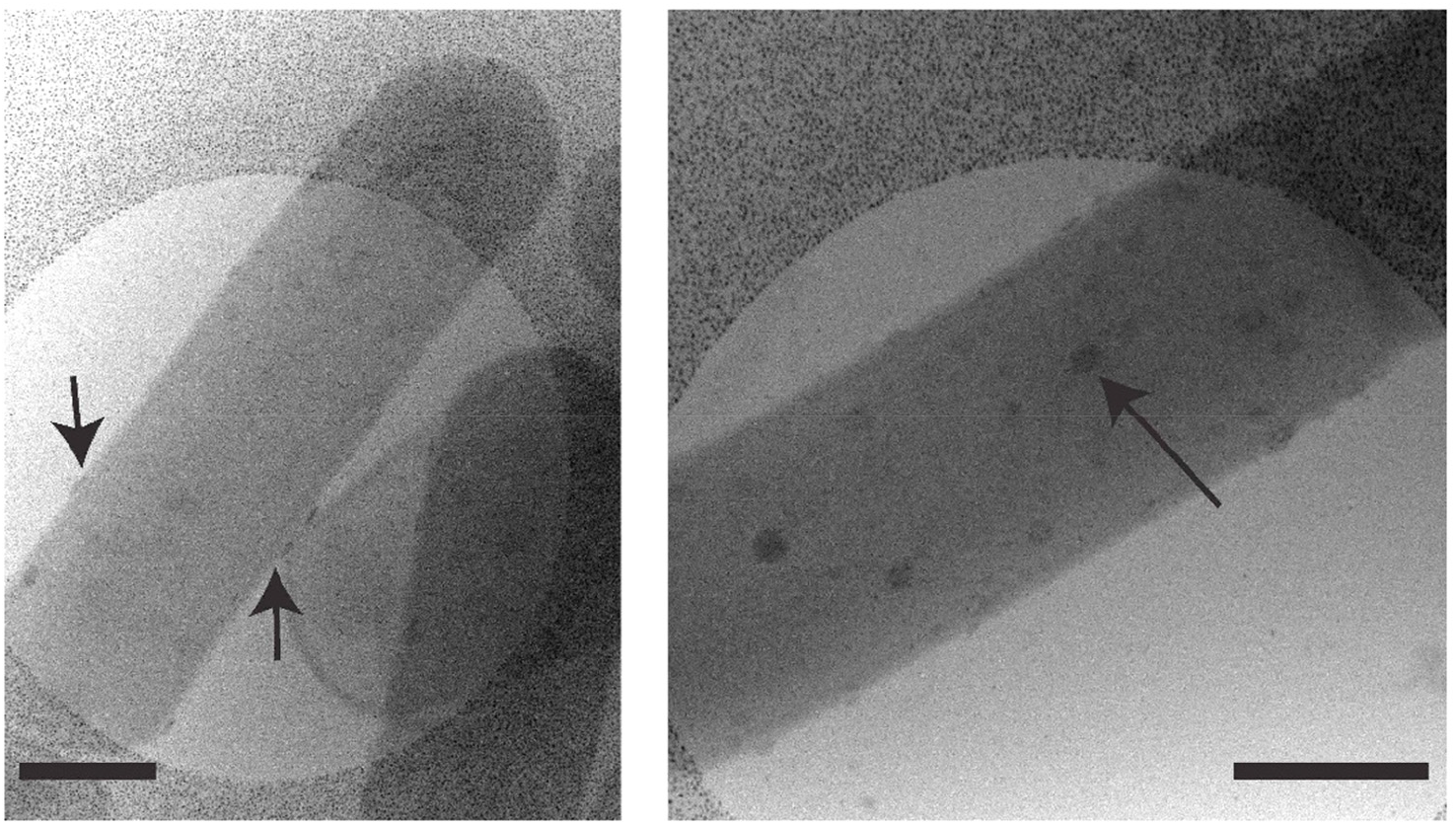
Cryo-STEM bright-field images of representative biofilm cells from a Δ*sigF* colony grown for 10 days on B4 agar. Black arrows point to aggregates with edge-on views (left panel) and top views (right panel). Scale bars 500 nm. A representative field imaged (out of n=30 cells), from 4 independent experiments is shown.

**Supplementary Figure 8.**
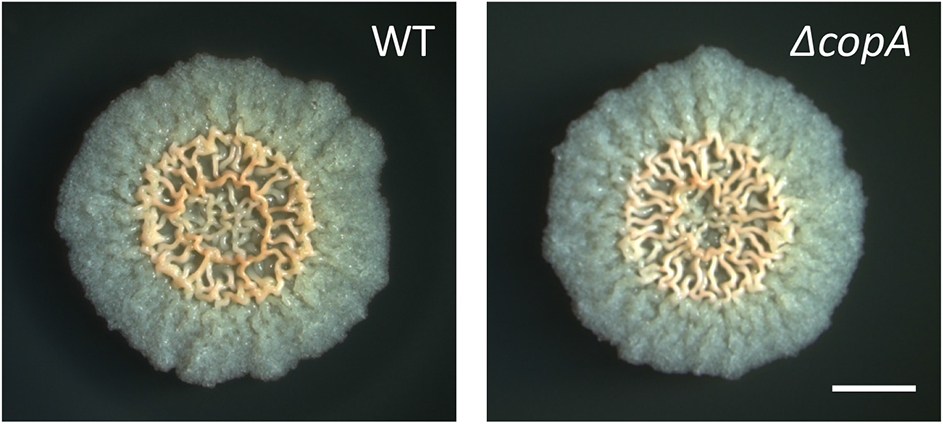
Light microscopy images of wild type *B. subtilis* (WT) and Δ*copA* mutant. Biofilm colonies were grown on B4-Ca^2+^ agar for 3 days at 30°C. Scale bar – 2 mm. The experiment was repeated 3 times, in a technical quadruplicate – and representative images are shown.

**Supplementary Figure 9.**
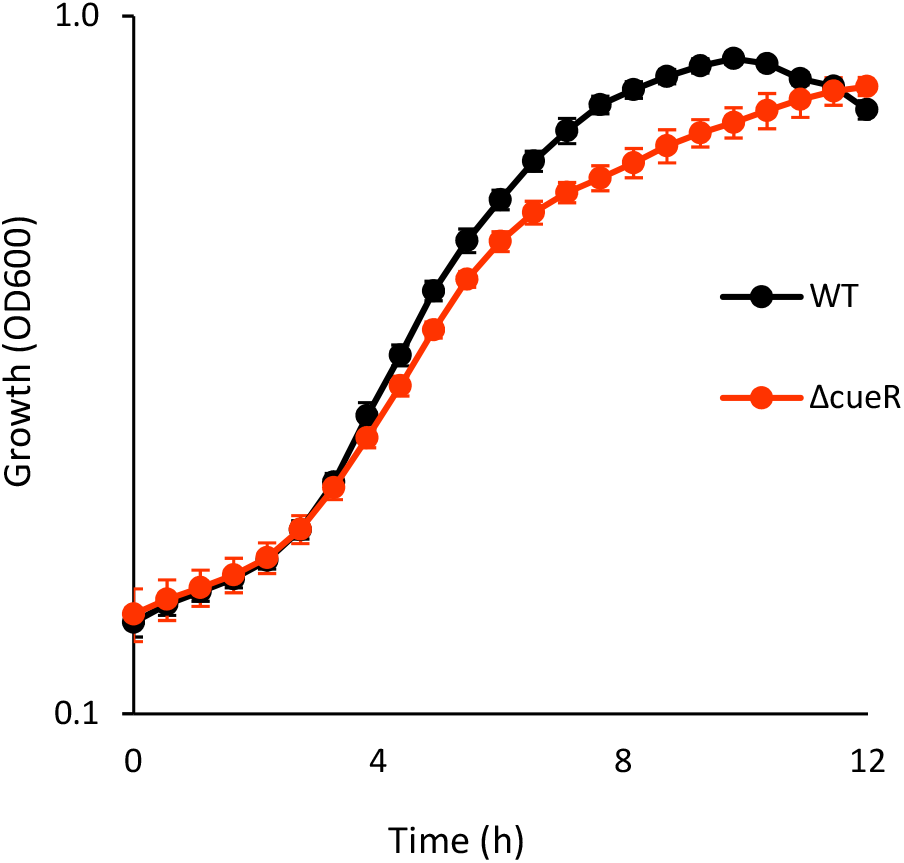
Planktonic growth assay. Wild type *B. subtilis* (ET) and *ΔcuerR* mutant were grown at 30°C with shaking in liquid LB, and growth was monitored by measuring OD600 in a microplate reader every 30 min. Results are averages of six wells, bars represent standard deviations. A representative experiment out of at least 3 independent experiments is shown.

**Supplementary Figure 10.**
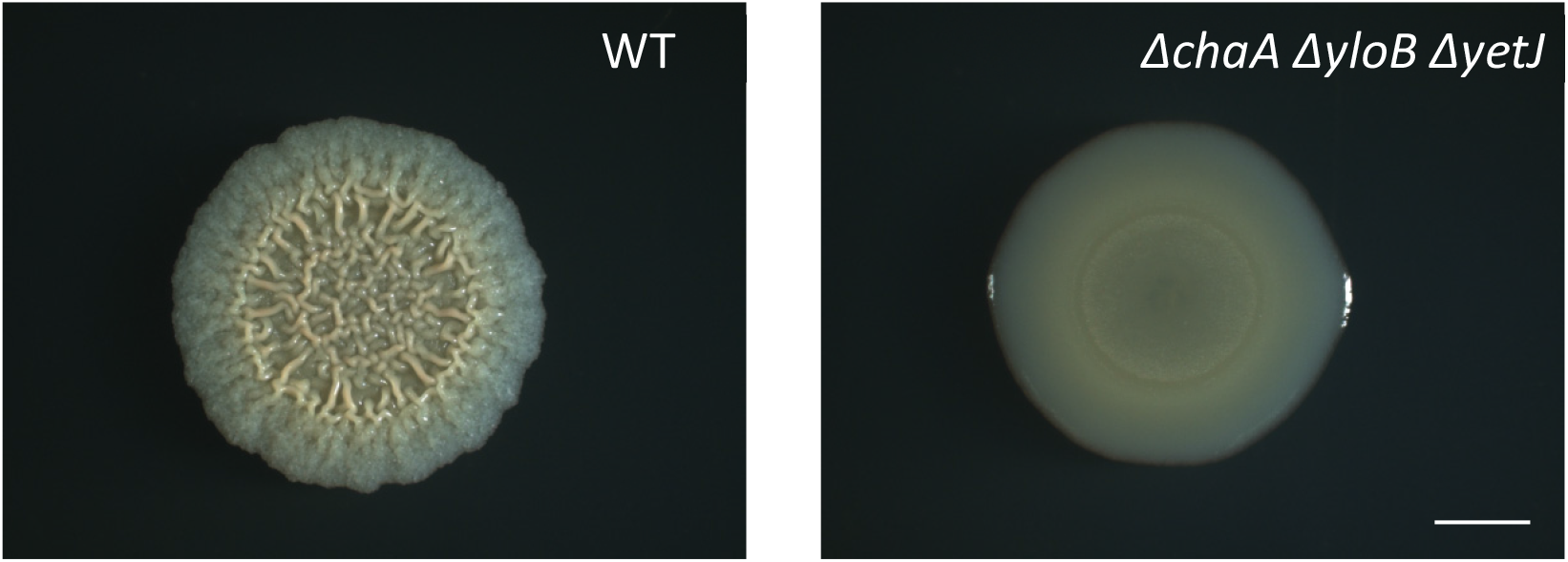
Light microscopy images of wild type (WT) and triple *ΔchaA ΔyloB ΔyetJ* mutant. Biofilm colonies grown on solid B4-Ca^2+^ agar for 3 days at 30°C. Scale bar – 2 mm. The experiment was repeated 3 times, in a technical quadruplicate – and representative images are shown.

**Supplementary Figure 11.**
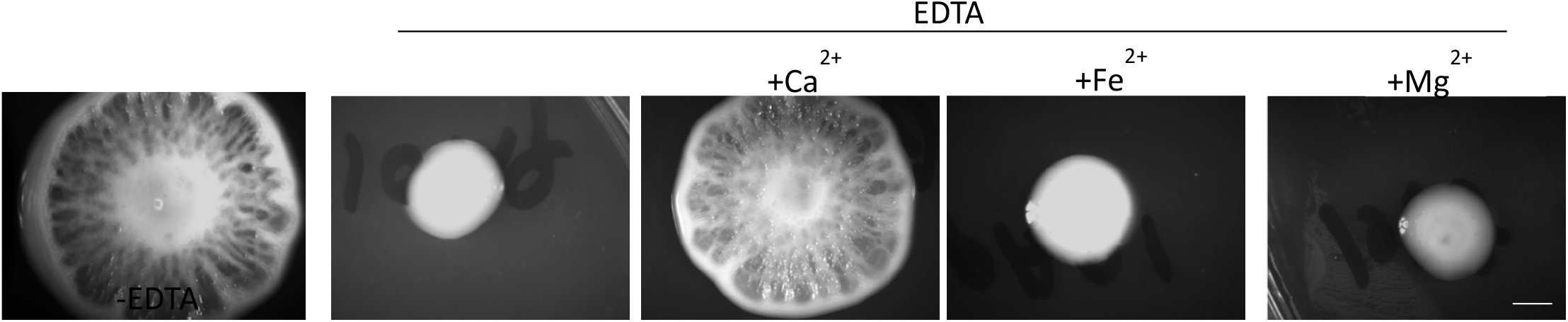
Light microscopy of biofilm colonies of *P. aeruginosa* PA01 supplemented with EDTA (0.1mg/ml), and divalent ions (1mM), as indicated. Colonies were grown on BHI solid medium for 4 days at 23°C. Scale bar – 5 mm. The experiment was repeated 3 times, in a technical 3 repeats – and representative images are shown.

**Supplementary figure 12:**
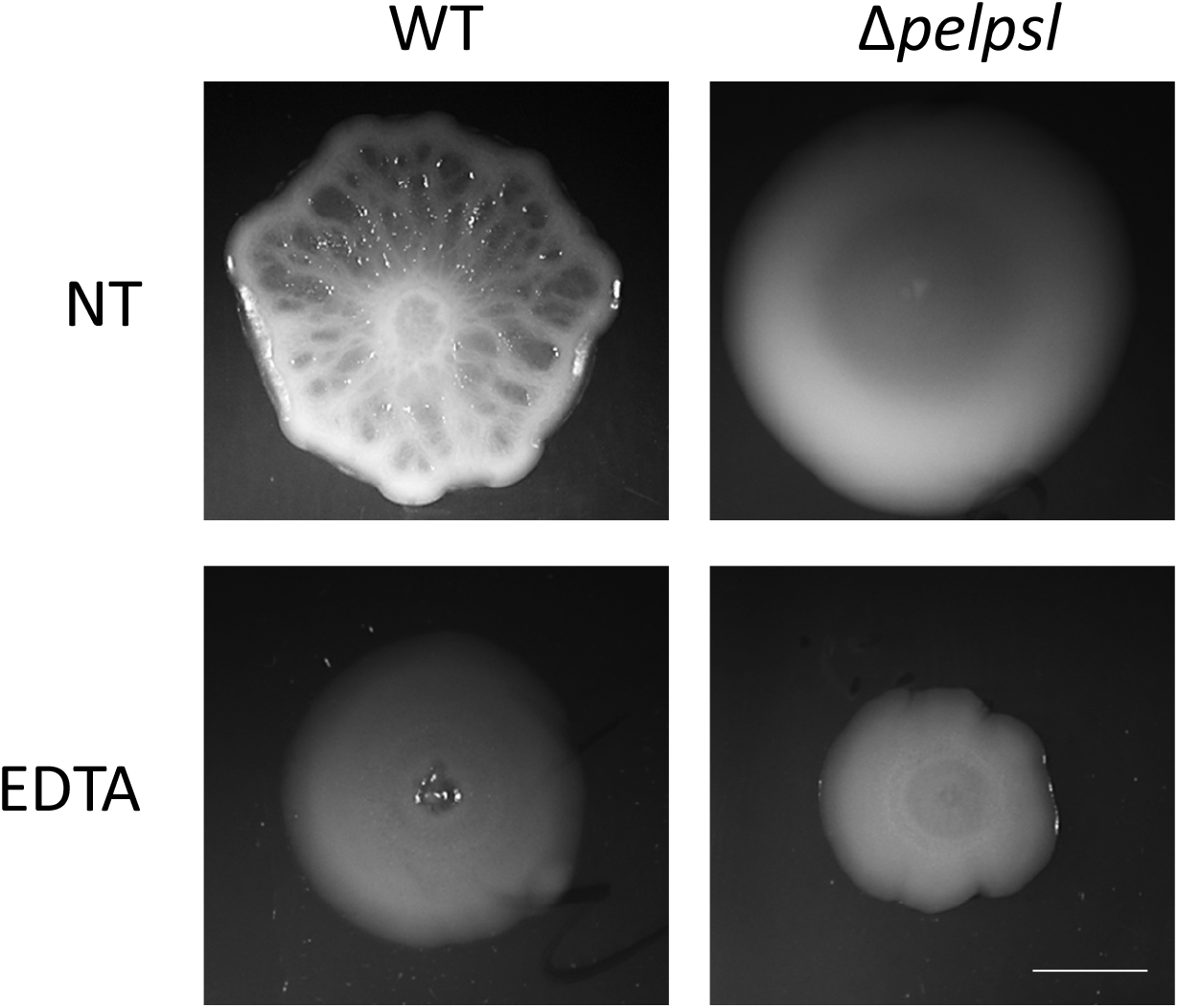
Light microscopy of biofilm colonies of *P. aeruginosa* PA01 either untreated or supplemented with EDTA (0.1mg/ml) as indicated. Colonies were grown on BHI solid medium for 4 days at 23°C. Scale bar = 5 mm. The experiment was repeated 2 times, in a technical 3 repeats – and representative images are shown.

**Supplementary Figure 13.**
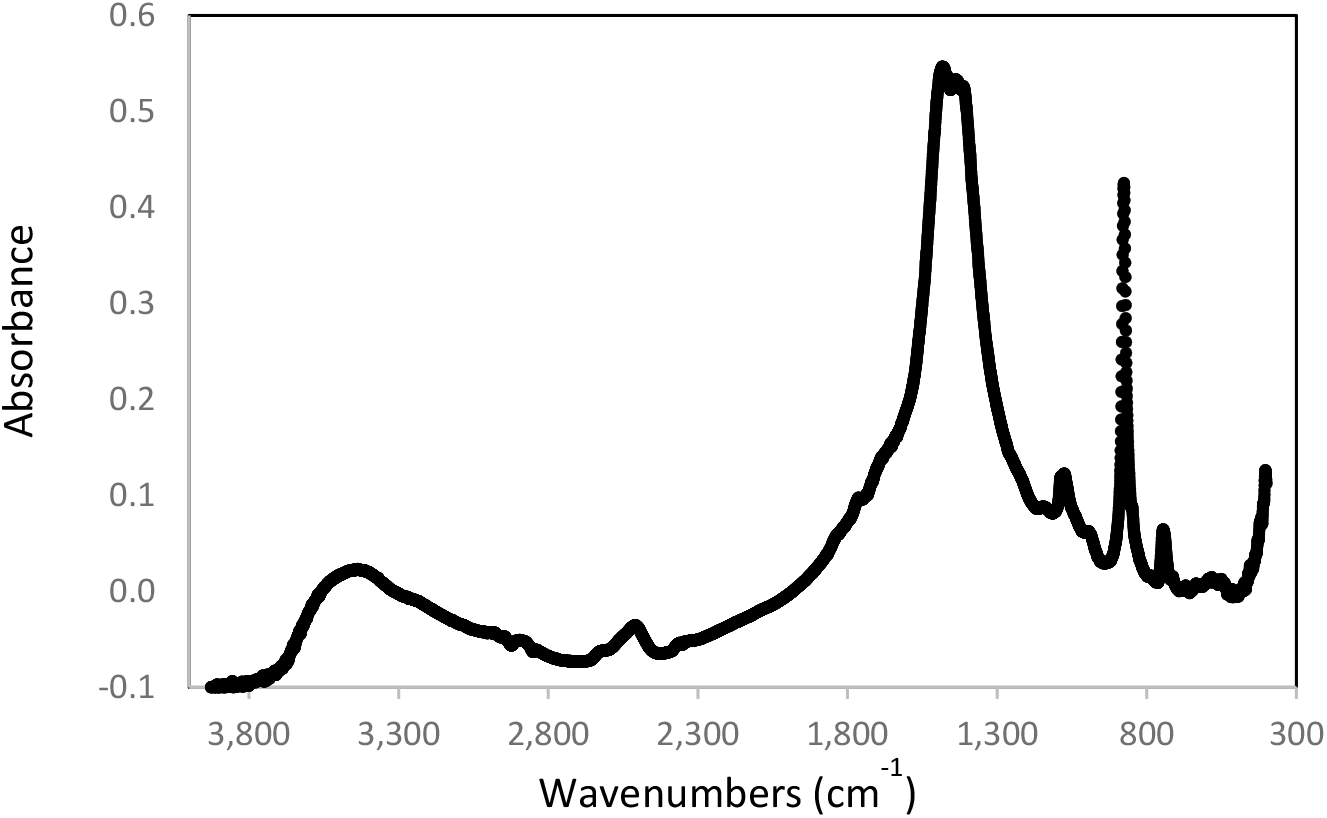
FTIR spectra of bleached *M. abscessus* biofilm colonies, displaying vibrations characteristic of varterite (crystalline calcium carbonate).

**Supplementary Figure 14.**
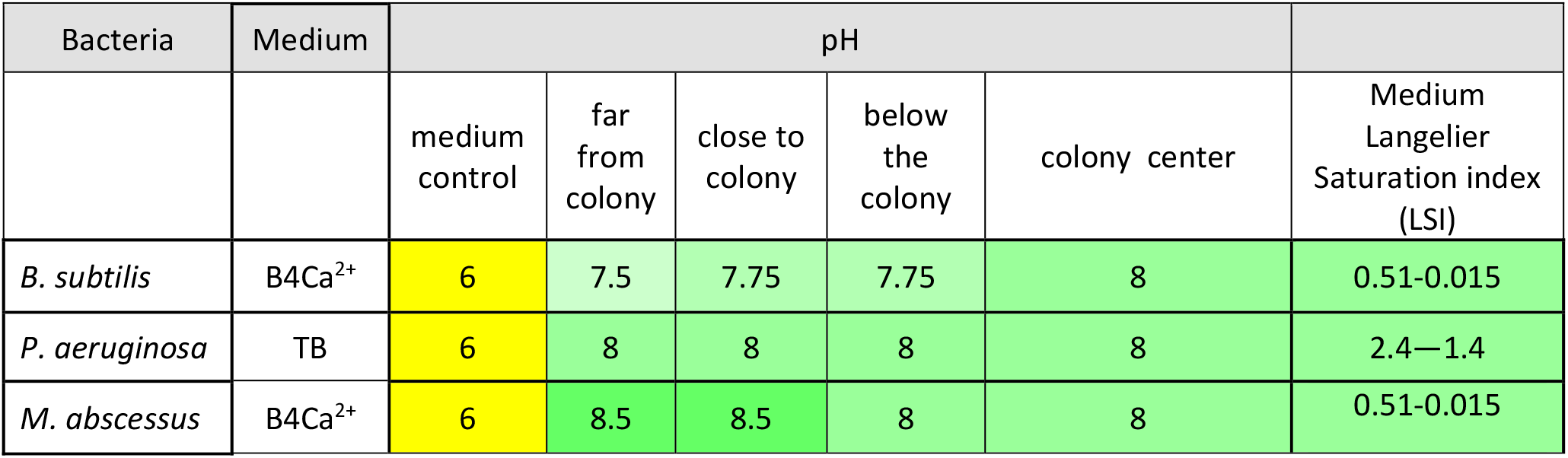
pH of colonies and the agar medium they were grown on was assessed using pH-indicator strips (MColorpHast™). The experiment was conducted at least twice for each medium, with at least 4 technical repeats. LSI calculation refers to the indicated growth media. ND – undetermined.

**Supplementary Figure 15.**
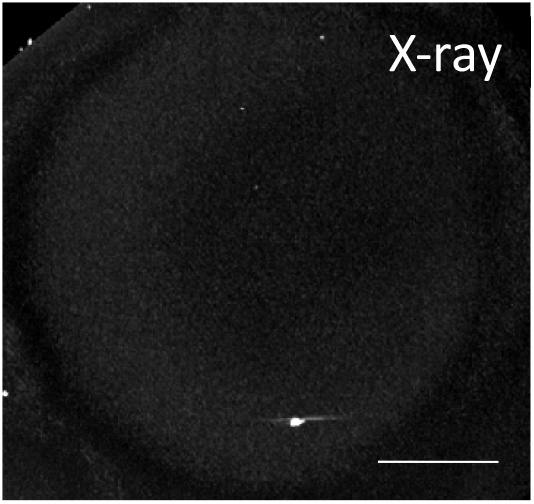
MicroCT-XRay image of a wild type *P. aeruginosa* PAO14 colony grown for 3 days at 23°C on TB agar containing SMV (0.01 mg/ml). The absence of visible bright areas indicates lack contrast, indicating no dense mineral (compare to the control colony shown in Fig. 3A). Scale bar – 2 mm. The experiment was repeated twice with similar results.

**Supplementary Figure 16.**
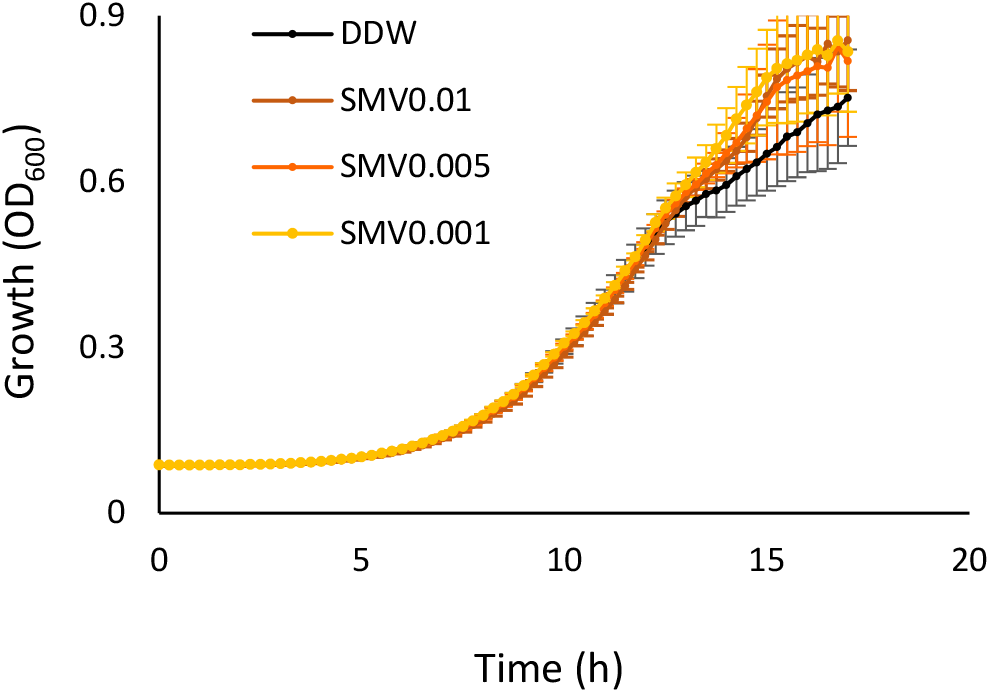
Planktonic growth assay. *P. aeruginosa* PA14 was grown at 37°C with shaking in liquid TB medium, supplemented with: 1.5 mg/ml DTNB, 1.75 mg/ml AHA, 2.5 mg/ml Diamox and 0.01, 0.05 and 0.01 mg/ml SMV. Growth was monitored by measuring OD600 in a microplate reader every 30 min. Results are averages of six wells, bars represent standard deviations. A representative experiment out of at least 3 independent experiments is shown.

**Supplementary Figure 17.**
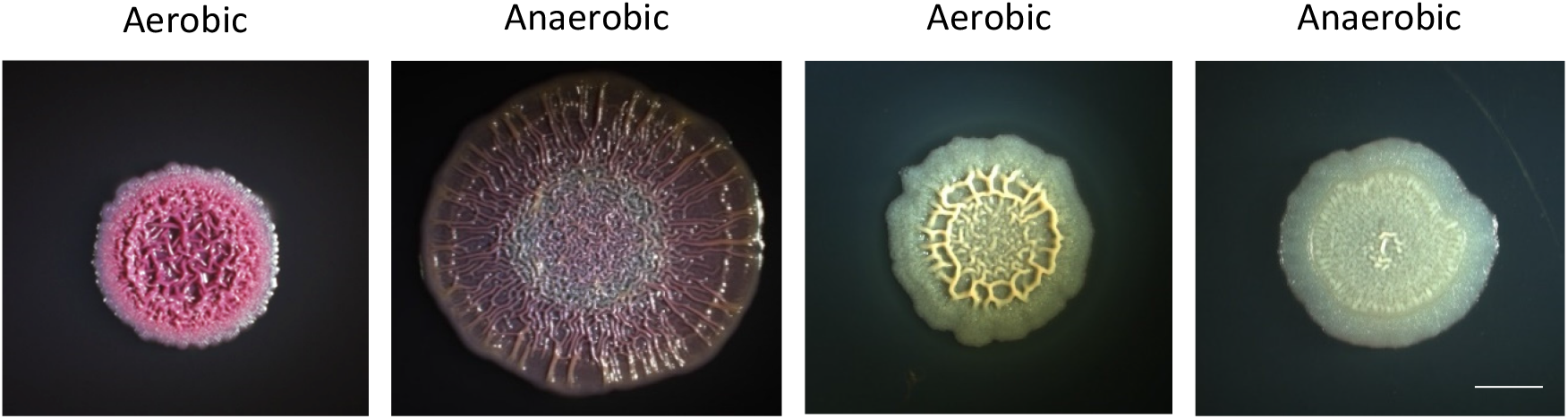
Light microscopy of biofilm colonies formed by *P. aeruginosa* PA14 (left) and *B. subtilis* (right). The biofilms were grown TB and B4-Ca^2+^ agar, respectively, with nitrate source (0.5% KN0^3^). Under aerobic conditions, CO_2_ is available from the atmosphere and also produces as a byproduct of aerobic respiration. Under anaerobic conditions of nitrate reduction (as used here - anaerobic growth chamber, a mix of H_2_ and N_2_ (5/95%), CO_2_ is absent from the atmosphere, and aerobic respiration (and subsequent CO_2_ emission) is not feasible due to the lack of oxygen. The experiment was repeated 4 times, in a technical 3 repeats– and representative images are shown.

**Supplementary Figure 18.**
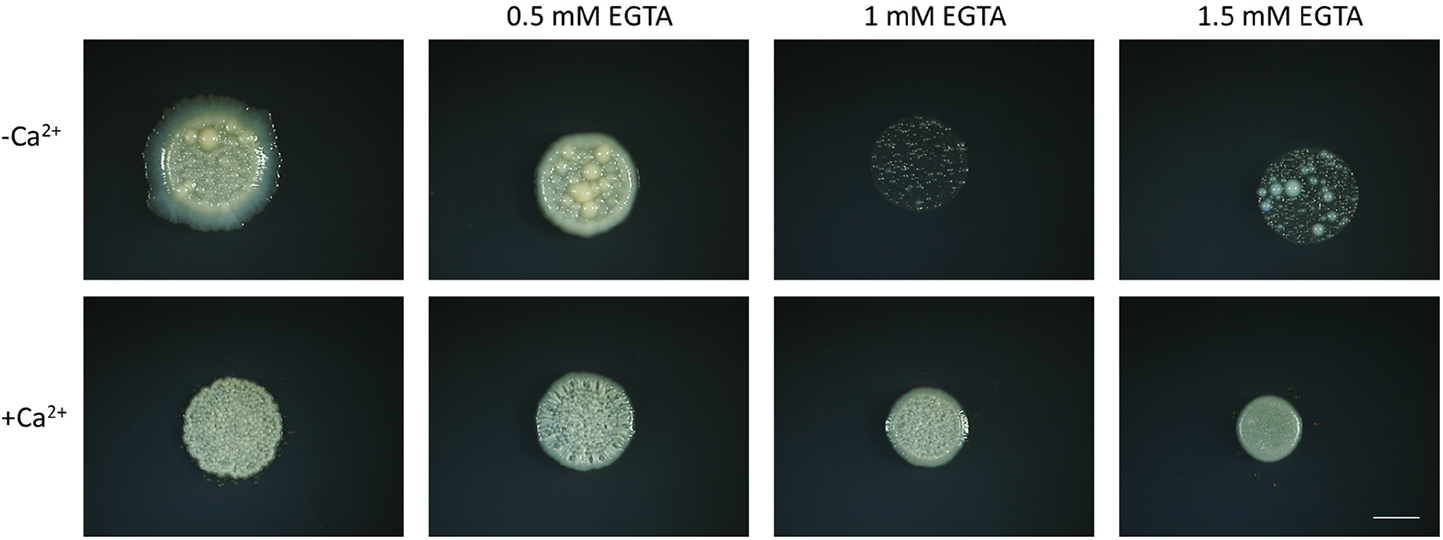
Light microscopy of 3-day-old biofilm colonies formed by *M. abscessus* on B4 agar supplemented with calcium acetate (0.25% v/v) and EGTA, as indicated. Scale bar = 5 mm. The experiment was repeated 3 times, in a technical 3 repeats– and representative images are shown.

**Supplementary Figure 19.**
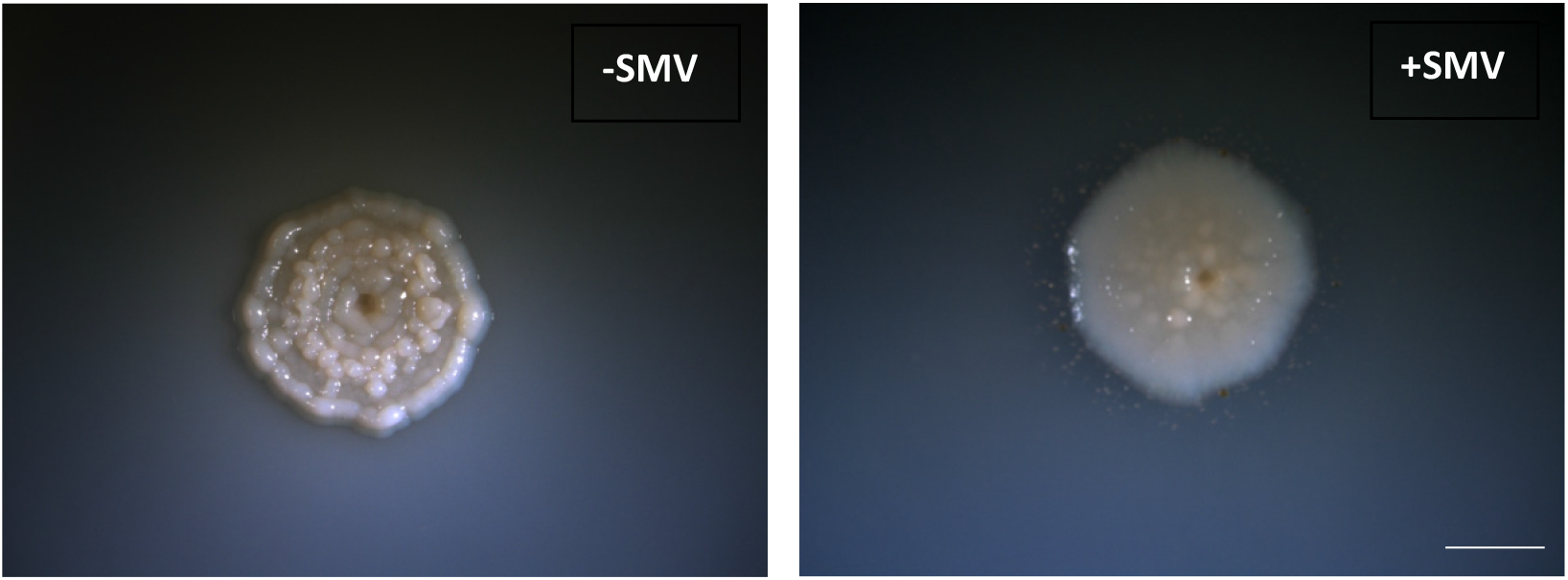
Light microscopy images of biofilm colonies formed by *M. abscessus*, supplemented with SMV (0.025mg/ml) as indicated. Colonies were grown on B4-Ca^2+^ medium for 5 days, at 37°C. Scale bar = 5 mm. The experiment was repeated 2 times, in a technical 3 repeats – and representative images are shown.

**Supplementary Figure 20.**
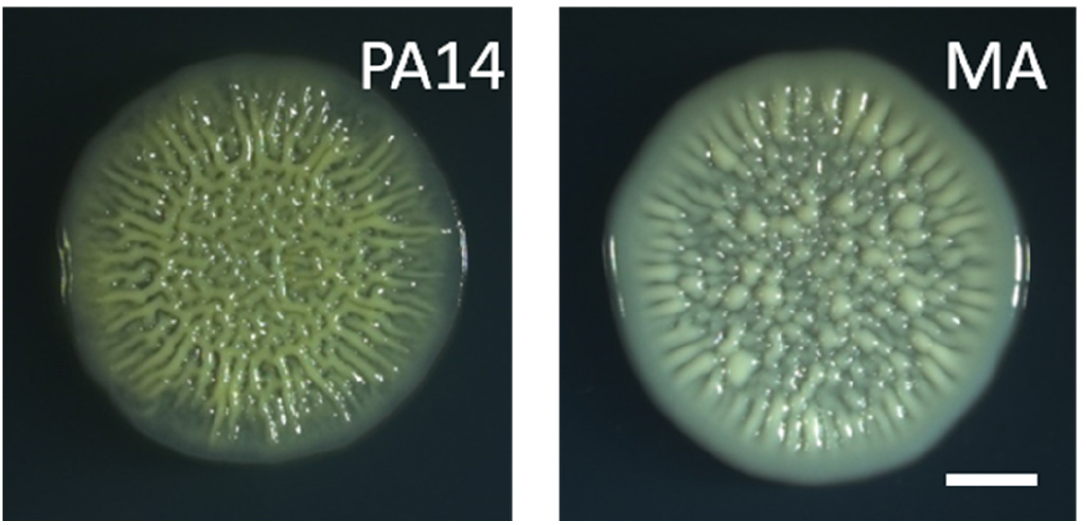
*P. aeruginosa* and *M. abscessus* biofilm colonies grown on solid SCFM medium for 4 days at 23°C (*P. aeruginosa*) and 30°C (*M. abscessus* and *S. aureus*). Scale bar – 2 mm. The experiment was repeated 3 times, in a technical 3 – and representative images are shown.

**Supplementary Figure 21.**
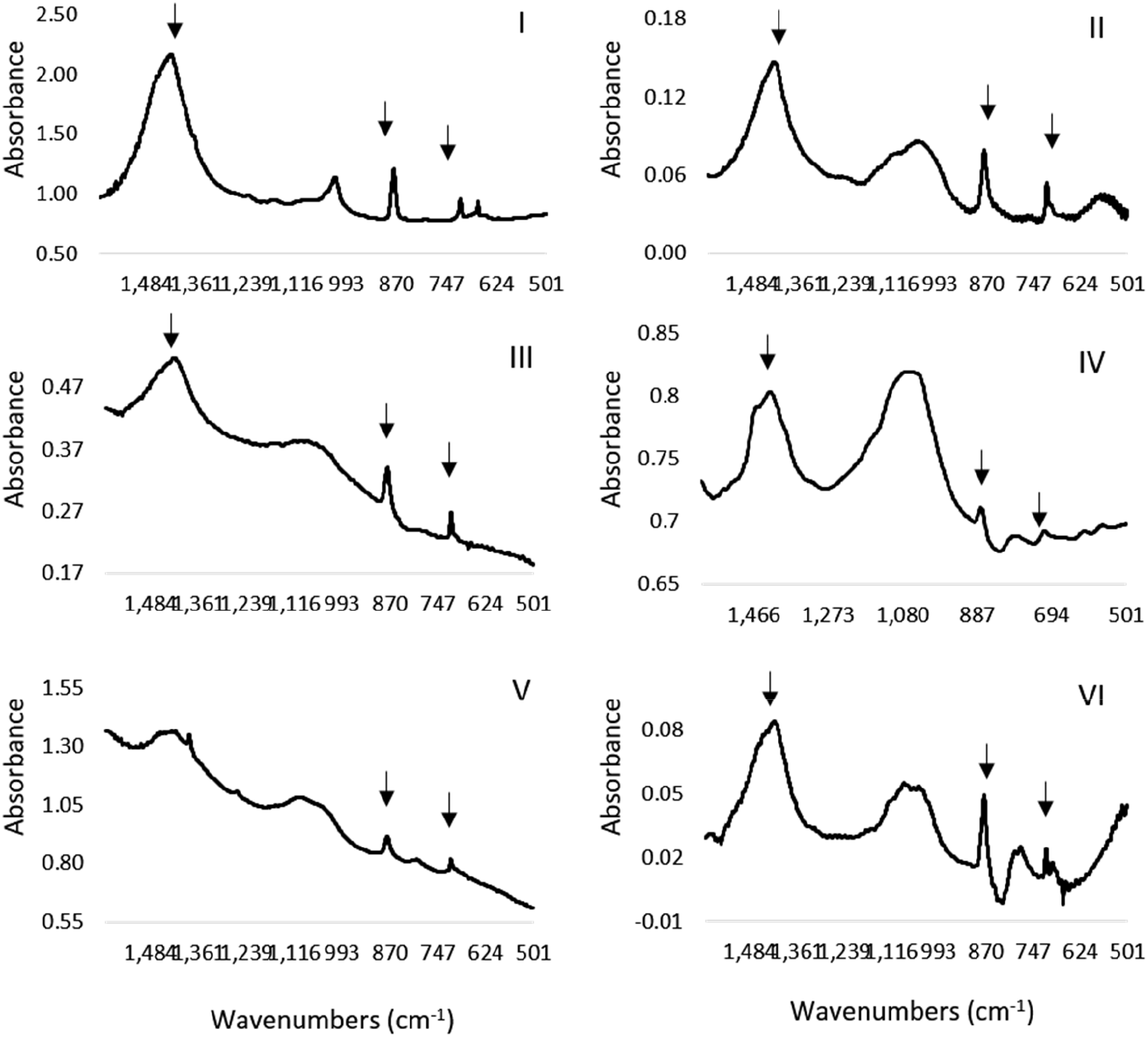
FTIR spectra of bleached sputum samples of *P. aeruginosa* positive CF patients (I-VI). Arrows indicate vibrations characteristic of calcite. For patient details, see Supplementary table 2.

**Supplementary Figure 22.**
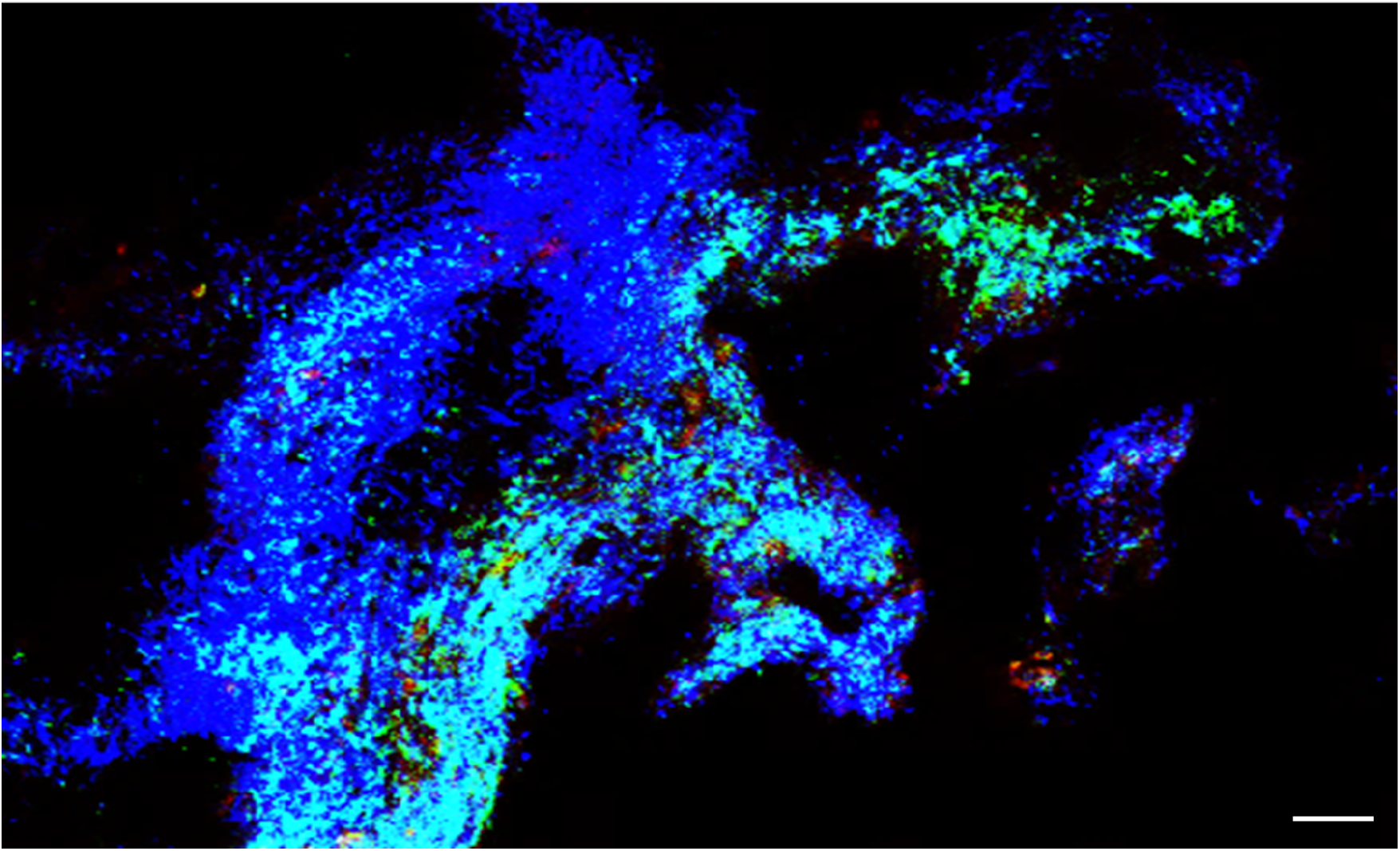
Confocal image of *ex-vivo* lung model. Lungs were harvested from one-month-old mice, cultured in DMEM containing carbenicillin 100 µg/ml, and infected with *P. aeruginosa* PA14 constitutively expressing GFP. Green – bacterial cells. Blue - nuclei of lung cells stained with DAPI. Scale bar – 50 µm. The experiment was repeated 3 times, in a technical 4 repeats – and representative image is shown.

**Supporting Figure 23.**
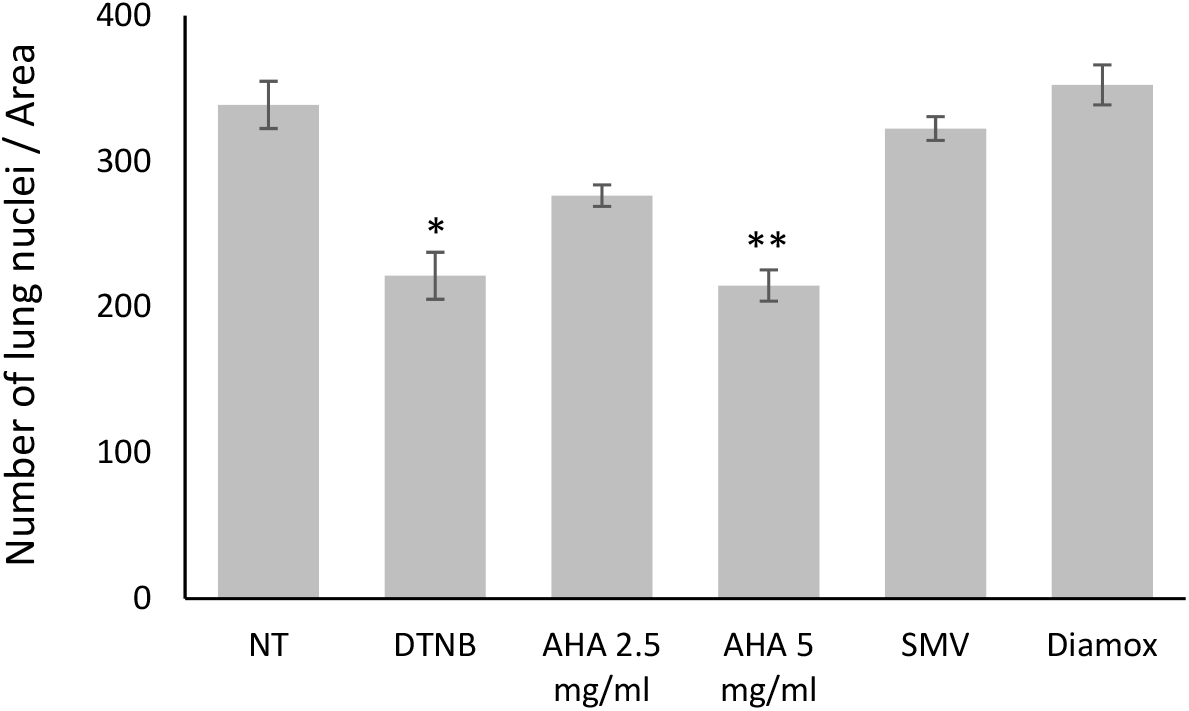
Quantification of lung viability. Lung tissue cultures were treated with biomineralization inhibitors DTNB (2 mg/ml), SMV (0.01 mg/ml), Diamox (1.1 mg/ml) and AHA, as indicated. ImageJ 1.51g software was used to automatically count lung cell nuclei in 4 randomly chosen fields for each treatment. P values, as determined by student’s t-test, are indicated (* pVal <0.01, ** pVal <0.001).

**Supporting Figure 24.**
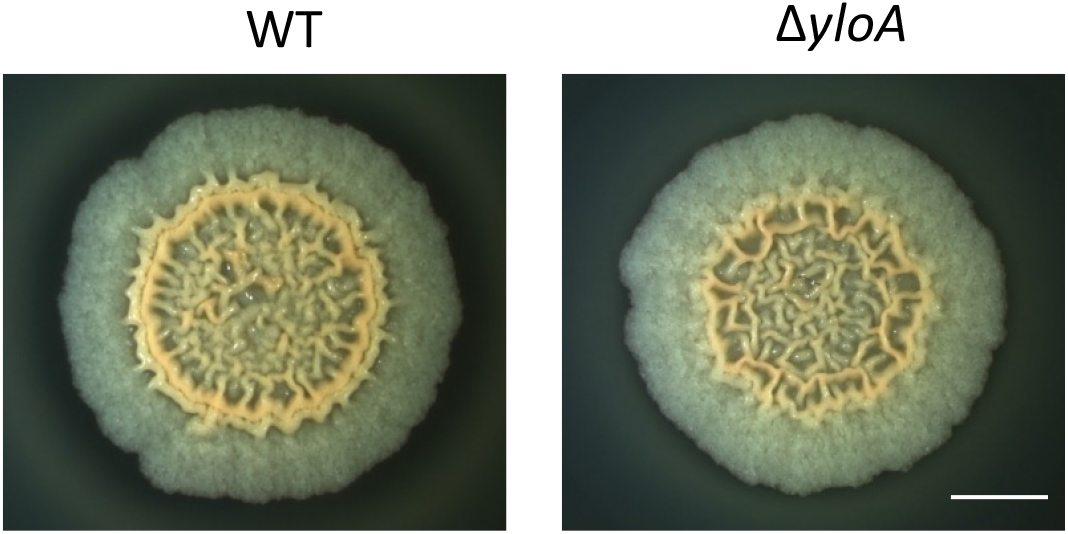
Light microscopy images of wild type (WT) and *ΔyloA* mutant. Biofilm colonies grown on solid B4-Ca^2+^ agar for 3 days at 30°C. Scale bar – 2 mm. The experiment was repeated 3 times, in a technical quadruplicate – and representative images are shown.

**Supporting Figure 25.**
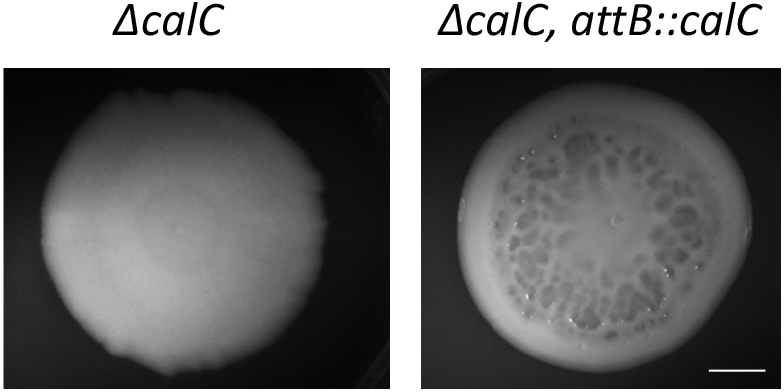
Light microscopy of biofilm colonies of *P. aeruginosa* PA01 indicated strains. Colonies were grown on BHI solid medium for 4 days at 23°C. Scale bar = 5 mm. The experiment was repeated 3 times, in a technical 2 or more repeats – and representative images are shown.

## Supplementary Files

**Supplementary File 1: Differentially expressed genes in *B. subtilis* NCIB 3610, + vs - calcium.**

**Supplementary File 2: Differentially expressed genes *B. subtilis* NCIB 3610, WT vs *cueR*.**

**Supplementary File 3: Patients information.** Clinical parameters of patients analyzed for presence of calcite in sputum samples.

## Supplementary Movies

**Supplementary Movie 1:** *B. subtilis* colony

**Supplementary Movie 2:** *P. aeruginosa* PA14 colony

**Supplementary Movie 3:** *M. abscessus* colony

The intact colonies were visualized under X-ray, and the obtained 2D images were used to generate high-resolution 3D image (see Keren-Paz *et al.,* 2018 for details). Rotation of 360° was taken around an axis perpendicular to the biofilm surface. Results are of a representative experiment out of three independent repeats.

## Supplementary Tables

**Supplementary Table 1:** **Strains used in this study**

| Genotype | Reference |
| --- | --- |
| *Bacillus subtilis* NCIB 3610 | Branda *et al* (2001) |
| $\Delta arsR$ (kan) | This work |
| $\Delta mntR$ (kan) | This work |
| $\Delta zur$ (kan) | This work |
| $\Delta cueR$ (kan) | This work |
| $\Delta copA$ (kan) | This work |
| $\Delta yetJ$ (spec) | This work |
| $\Delta chaA$ (kan) | This work |
| $\Delta yloB$ (mls) | This work, the lack of polar effect was confirmed by the deletion of *yloA* (Figure S24) |
| $\Delta chaA$ (kan) $\Delta yetJ$ (spec) | This work |
| $\Delta chaA$ (kan) $\Delta yetJ$ (spec) $\Delta yloB$ (mls) | This work |
| *Pseudomonas aeruginosa* PA14 | Lab collection |
| *Pseudomonas aeruginosa* PA14, Ppuc-GFP (carb) | <sup>1</sup> |
| *Pseudomonas aeruginosa* PA01 | Lab collection |
| *Pseudomonas aeruginosa* PA01 $\Delta psCA1$ | <sup>1</sup> |
| *Pseudomonas aeruginosa* PA01 $\Delta psCA1-2$ | <sup>1</sup> |
| *Pseudomonas aeruginosa* PA01 $\Delta psCA1-2-3$ | <sup>1</sup> |
| *Pseudomonas aeruginosa* PA01 $\Delta calC$ | This work, This work, the lack of polar effect was confirmed by complementation of *calC* from an ectopic chromosomal loci (Figure S25), $\Delta calC$ attB::*calC* |
| *Mycobacterium abscessus* ATCC 19977 | A kind gift from Daniel Barkan |

- 1 Lotlikar, S. R. *et al.* *Pseudomonas aeruginosa* beta-carbonic anhydrase, psCA1, is required for calcium deposition and contributes to virulence. *Cell Calcium* **84**, 102080, doi:10.1016/j.ceca.2019.102080 (2019).

**Supplementary Table 2:**
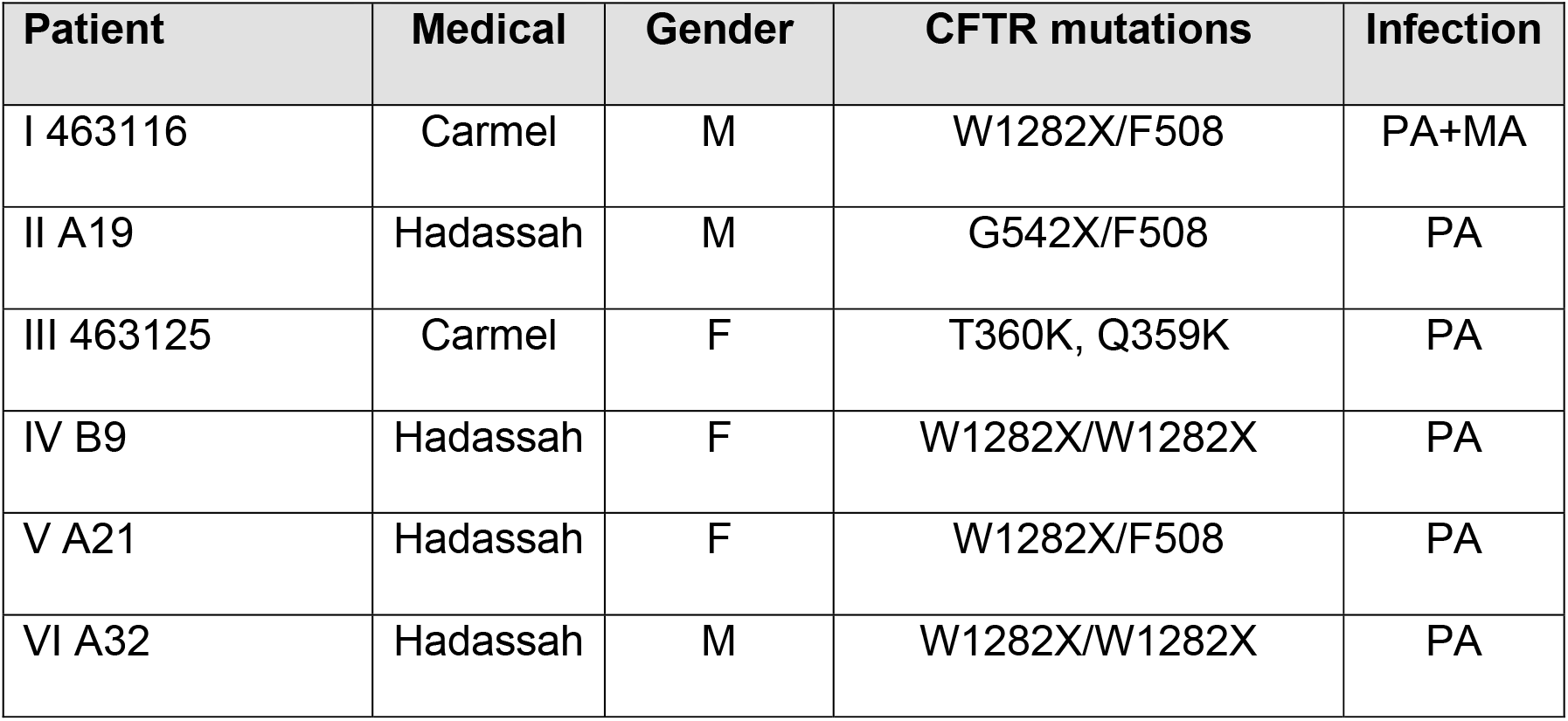

The details of patients included Supplementary Figure 15 and their respected panels are indicated.

